# Structural Composition Enables Very Fast Learning

**DOI:** 10.64898/2026.07.14.738142

**Authors:** Reidar Riveland, Alex Pouget, Peter Latham

**Affiliations:** Gatsby Computational Neuroscience Unit, University College London, London, UK; Department of Basic Neuroscience, Université de Genève, Geneva, Switzerland

## Abstract

There is a gap between neuroscientific theories of learning and the speed of learning observed in many experiments. Since the Cognitive Revolution of the 1950s, compositionality has played a central role in efforts to bridge this gap. Roughly, a compositional system is one where distinct modules are combined according to a set of rules in order to accomplish complex tasks. Recently, significant progress has been made in understanding the emergence of modules in both biological and artificial neural systems. How, and under what conditions, the rules of module recombination are represented in these systems remains an open question. Here we present a neural model that can leverage these rules to dramatically speed up learning. We first show that when faced with multiple tasks which share subcomponents, models learn a low-dimensional representation that captures how subcomponents are reused across the task set. These low-dimensional spaces encode the structure that governs how modules should be recombined. Restricting learning to these subspaces greatly reduces the amount of experience needed to acquire a novel task, even when learning from reinforcement on single trials. In some cases, we can leverage the geometric regularities of these representations to reduce learning to a form of hypothesis testing over a small set of discrete points. Finally, we use this theory to model both behavioral and neural data from non-human primates performing a compositional task, and show that key features in this data are consistent with a model in which exploration during learning is restricted to these low-dimensional spaces. Overall, this work shows that the advantages of modularity in neural systems can be greatly improved upon when models represent the structure of module reuse. Both these features working in tandem lead to learning on timescales similar to biological intelligences, and hence provide a model for how such fast, adaptable behavior can emerge from systems of neurons.

## Introduction

One of the oldest and most successful ideas in neuroscience is that learning occurs because of slow changes in synaptic weights [1–4]. Generally, changing weights to adapt to new circumstances predicts a gradual acquisition of skills. By contrast, learning observed in many experiments can occur in a relatively small number of trials or involve sudden insights [5, 6]. How to bridge the divide between slow weight updates and the rapid adaptability of biological intelligence is a core problem of brain and cognitive science [7–9]. In the middle of the 20th century, computational cognitive science proposed a solution: compositionality. A compositional system is one where distinct modules are combined according to a set of rules in order to accomplish a complex task. Originally, a framework for understanding the expressivity of natural language [10], compositionality soon spread beyond language to become a more general principle used to explain cognitive phenomena in domains as diverse as navigation [11, 12], vision [13–15], motor planning [16], and reasoning [17, 18].

For neuroscientific theories of learning, the appeal of compositionality left open the question of how such a scheme might be implemented in neural systems. Early neural network models appeared incapable of implementing composition, sparking debate over whether neural compositionality was possible in principle [19, 20]. However, converging evidence from advances in neural network modeling and recordings taken during increasingly sophisticated animal behaviors suggests that compositional computations can indeed emerge in networks of neurons [21–29]. In a prominent line of work studying recurrent neural networks (RNNs) performing a variety of cognitive tasks, the authors found that networks reused modules across tasks with similar demands (e.g. they reuse the same ring attractor across tasks that required memorizing a circular variable) [21]. Importantly, after the network developed these modules during pre-training, they could be recombined in new ways to perform novel tasks without changing the recurrent weights. Instead, networks simply needed to learn a new input which configured the RNN to recruit proper recombination of modules for the novel task. We call this input, which is typically a high-dimensional vector, a *task embedding vector*.

The central claim of this work is that if subjects take full advantage of regularities in how modules are used across the task set, learning can be extremely fast – often requiring only single digit number of trials. This is possible because, in compositional settings, there are a small number of dimensions in the space of task embeddings which fully describe how modules are reused, and those dimensions effectively control the entire system. Restricting learning to these subspaces results in very fast acquisition of new tasks. In effect, this is a neural implementation of the rules of module recombination – combinations that are valid according to the experience of pretraining all reside within these subspaces.

In what follows, we first show empirically that task representations in multitasking RNNs encode the compositional structure needed to make rapid learning possible in principle. To understand the emergence of this structure, we analyze a feedforward network trained on analogous compositional task sets. In this setting, we precisely characterize how the organization of the task set relates to structure in learned task embeddings, and describe the conditions required for this correspondence to hold. We conjecture that the mechanism we find in the feedforward network applies to recurrent networks as well, and validate this across many task sets learned by RNNs. We then show how systems can use this structure in task embeddings to achieve rapid learning. Finally, we demonstrate that features of behavioral and neural data recorded from non-human primates performing a compositional task are consistent with learning that is guided by this structure.

Crucially, these results hold for systems without architectural choices designed to capture compositional structure. Instead, this structure emerges naturally from performing tasks with overlapping demands. At a high level, we see improvement in the speed of learning because our system doesn’t just recombine modularized computations. Rather, by learning in the relevant subspace it also implicitly follows the rules – or the structure – of recombination observed during training. Joining the complementary ideas of modules and structure moves us closer to realizing powerful forms of compositionality in neural systems, and allows us to model fast and flexible learning observed in biological intelligences.

## Results

### Modularity and task set structure in multitasking recurrent networks as a path to modeling rapid learning

We begin with a concrete example. Consider a set of decision-making tasks that consist of multiple interchangeable parts, chosen to closely resemble those used in the experimental literature [29– 32]. In these tasks, participants receive sensory input in two input modalities (Mod1 and Mod2) representing, for example, audition and vision. On every trial the input, which appears in both modalities, consists of a strong and a weak signal, placed at random points on a circle (Fig. 1a, left panel). The trial unfolds over time, and includes a fixation period, followed by stimulus presentation, and then a response period. The demands of each task are defined by three binary axes. The first determines the input modality to attend to (Mod1 versus Mod2; e.g., pay attention to vision and ignore audition). The second determines whether to respond to the strongest or the weakest stimuli presented in that modality (Strong vs. Weak). The third determines the proper method of response (Resp1 vs. Resp2; e.g., saccade vs. joystick). A schematic of the stimulus and response structure for example ‘Mod1-Strong-Resp1’ and ‘Mod2-Weak-Resp2’ trials are shown in Fig. 1a (the time dimension of the task in this figure is omitted for easy visualization; see Methods for full details). In total 8 tasks can be built out of the 6 task components (Mod1 vs. Mod2; Strong vs. Weak; Resp1 vs. Resp2). Crucially, the sensory inputs by themselves don’t allow participants to infer the task demands. Therefore, a task cue is provided. In simulations these are encoded as one-hot vectors.

**Figure 1:**
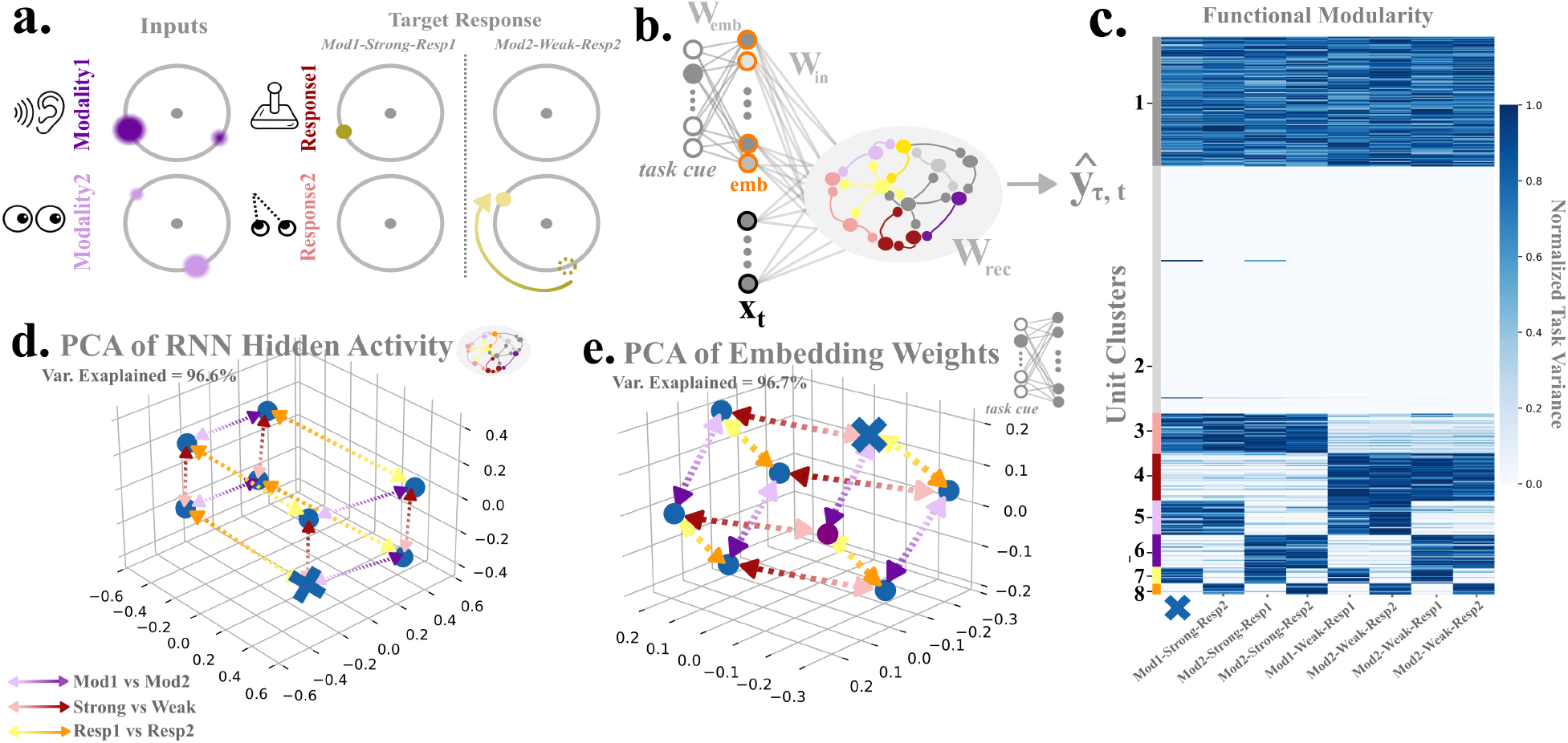
Modularity and task set structure in multitasking recurrent networks. **a)** Input and target output structure for two tasks in our decision-making (DM) task set. In the Mod1-Strong-Resp1 task, the participant must attend to stimuli only in the first sensory modality (e.g. audition), respond to the strongest direction, and respond using the first response modality (e.g. joystick). In the Mod2-Weak-Resp2 task, one must attend only to the second modality (e.g. vision), respond to the weakest direction, and respond using the second response modality (e.g. saccade). **b)** Network architecture. The task demands are specified by a one-hot task cue, which linearly activates a hidden layer via a set of weights, **W**_emb_, to obtain a task embedding vector of activity for task *τ*, denoted **emb**_τ_(orange units). The task embedding and sensory input, **x**, are passed to the RNN via a set of input weights **W**_in_. Recurrent units are color coded to reflect the fact that after training they modularize computations. **c)** Task variance heatmap across recurrent units. Each column shows the extent to which individual neurons vary across different trial conditions of each task. Clustering reveals modular unit subpopulations responsible for computing different subcomponents of the task set. **d)** PCA of recurrent activity directly prior to stimulus presentation when input is driven purely by task embeddings. Points show averages over many trials. For each task activity lies along common set of three axes which each correspond to one of the three axes of the task set, Mod1 vs. Mod2, Strong vs. Weak, and Resp1 vs. Resp2 **e)** Same as **d** but for embedding weights **W**_emb_.

Although the 8 one-hot vectors (which are by definition orthogonal) cannot encode the three binary axes that organize the task set, it is nonetheless possible to learn the underlying organization in model weights. Indeed, the reuse of subcomponents according to a latent structure is typical of learning in these compositional settings across species [29, 33, 34]. Humans in particular have shown a remarkable aptitude for inferring underlying task rules efficiently, often in only a handful of trials [35, 36]. Given some experience with a subset of the tasks described above, we’d expect humans to learn a held out task in very few trials [37–39]. The primary goal of this paper is to describe the kind of neural architecture and learning algorithm that could account for the incredible speed with which biological organisms acquire tasks in such settings. We start by showing empirically that RNNs trained on the above task develop modular structure and some representation of how modules are reused across tasks. We then focus on the simpler case of bi-linear feedforward networks where we can characterize this structure precisely and outline conditions where it is guaranteed to emerge. Empirically, we then show across a variety of different kinds of task sets with different compositional organizations that the same predicted structure emerges in RNNs. These representations allow the systems to vastly constrain learning, resulting in very rapid acquisitions of novel tasks. We then turn to neural data recorded during compositional tasks and show that important features of this data are compatible with constraining learning in this manner.

We begin by training a recurrent neural network to perform the tasks described above. A schematic of the network architecture is shown in Fig. 1b; it differs from past work [21, 25] in that a single layer of weights, **W**_emb_, processes the task cue into a task embedding vector. This representation is fed into the recurrent neural network along with sensory input, **x**_*t*_, via input weights **W**_in_.

Previous work [21, 25] has shown that training on compositional task sets leads to modular activity, in that the variability of a neuron in a given task is determined by which task components are active on the task. For instance, one neuron might exhibit variability only when Mod1 is relevant; another only when the subject should pay attention to the Weak component of the stimulus, and so on (this is indicated schematically by color coded units in Fig. 1b). To quantify this, we trained a network on 7 of the 8 tasks, with the Mod1-Strong-Resp1 task held out. Following past work, we then measured how much each unit in the RNN varied across trials in each task [25]. This roughly captures the degree to which each unit is contributing to computations (see Methods, RNN Modularity Analysis). According to this measure, units tend to cluster by task component (Fig. 1c). For example, units in cluster 3 have particularly high variance on all tasks that require the Strong component, but don’t vary across tasks that require the Weak component, suggesting these neurons support the responses to the strongest displayed direction. These clusters support modularity in the sense that, all other activations being equal, the network can swap between a Strong and a Weak task response by “switching on” cluster 4 variability and “switching off” cluster 3 variability.

The modularity exhibited by these systems suggests that to learn the held out task, the model only needs to activate the proper combination of modules rather than retrain weights from scratch. Past work accomplished this by freezing the recurrent weights (as these are the weights that implement modular computations), input weights, and output weights, and updating only the activity of the task embedding vector, which we denote 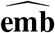. We use the notation 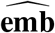 to denote task embedding activity learned for a novel task, and keep the *τ* subscript in **emb**_*τ*_ (Fig. 1c. orange) to denote task embedding activity for a task *τ* in the pre-training set. These are obtained by passing a one-hot vector of activity through embedding weights **W**_emb_. We call this process of learning 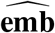 modular composition because optimization in the full task embedding space mainly takes advantage of the modularity in RNN computations. Though input, output and recurrent weights are frozen, this remains a high-dimensional optimization problem (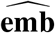 itself is high-dimensional). Past work has shown even with a supervised signal, learning 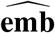 can take 100s of trials [22]. It is therefore unlikely that modular learning by itself can account for rapid acquisition of a new task, especially when learning from reinforcement on single trials.

Examination of the structure of RNN computations suggests an alternative way of choosing task embeddings that recruit the proper set of modules. Previous studies have noted that directly prior to stimulus onset the recurrent states of the RNN represent relevant dimensions of the task set [22, 25]. A visualization of this is shown in Fig. 1d, where the recurrent activity is organized along consistent Mod1 vs. Mod2, Strong vs. Weak, and Resp1 vs. Resp2 axes. This results in a 3-dimensional parallelogram geometry where each task resides at one of the vertices of this shape. It’s worth noting that representations which factorize task components along consistent axes in neural space have been consistently observed in animals and humans performing similar tasks [33, 34, 40]. Interestingly, we also observe this structure in the set of task embedding weights **W**_emb_, shown in Fig. 1e. This is the crucial fact that can, in principle, allow us to choose an embedding for the held out task quickly. Because of the geometric regularities in trained task embeddings, we can select an additional task embedding which we know lies on the Mod1, Strong, and Resp1 coordinates of this space (denoted by a blue cross in Fig. 1e.). Feeding this point in embedding space to the trained RNN leads to the activity denoted by the blue cross in Fig. 1d, recruits the set of modules in the first column of Fig. 1c, and leads to high performance on the held out task. This suggests that the embedding for the unseen task is closely related to embeddings for trained tasks, and knowing this can potentially constrain and therefore speed learning.

In the rest of the paper we’ll describe how such fast learning is possible, beginning with an analysis the structure of task embedding weights in compositional settings. In general, it is difficult to get an intimate mathematical understanding of nonlinear RNNs. Instead of analyzing RNNs directly, we hypothesize that the structure observed in **W**_emb_ (Fig. 1e) is primarily driven by structure in the task set, rather than recurrent dynamics. Under this hypothesis, our aim is to gain insight into the behavior of RNNs by studying similarly organized tasks in simple, analytically tractable, settings. In the following section we consider a bi-linear system with compositional structure that is analogous to the example decision-making task set considered above, and use this to demonstrate how the compositional structure of the task set is encoded in learned weights more generally. In later sections, we’ll leverage the insights gained about the structure of learned weights to build algorithms that use the structure of embedding weights to learn new tasks very rapidly.

### Two layer bi-linear networks provably represent compositional structure

We consider a set of *T* tasks, indexed by *τ*, which map input, **x** to the target output for the given task **y**_*τ*_ . The overall goal is to learn a function that takes inputs and task information to target outputs across different tasks,

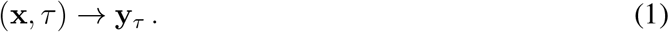

To make this problem analytically tractable, we ignore the temporal aspect of the task, and assume that the targets, **y**_*τ*_, are linear in **x**,

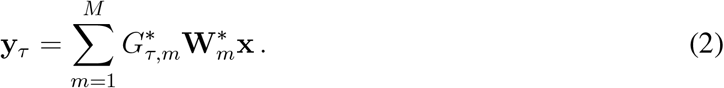

where **G**^∗^ is a *T* × *M* matrix that determines the extent to which modules contribute to any given task (*M* denotes the total number of modules; this is similar to mixture-of-experts [41] where here individual experts are mixed according to **G**^∗^ to produce training data). The network structure of this equation is illustrated in Fig. 2a.

**Figure 2:**
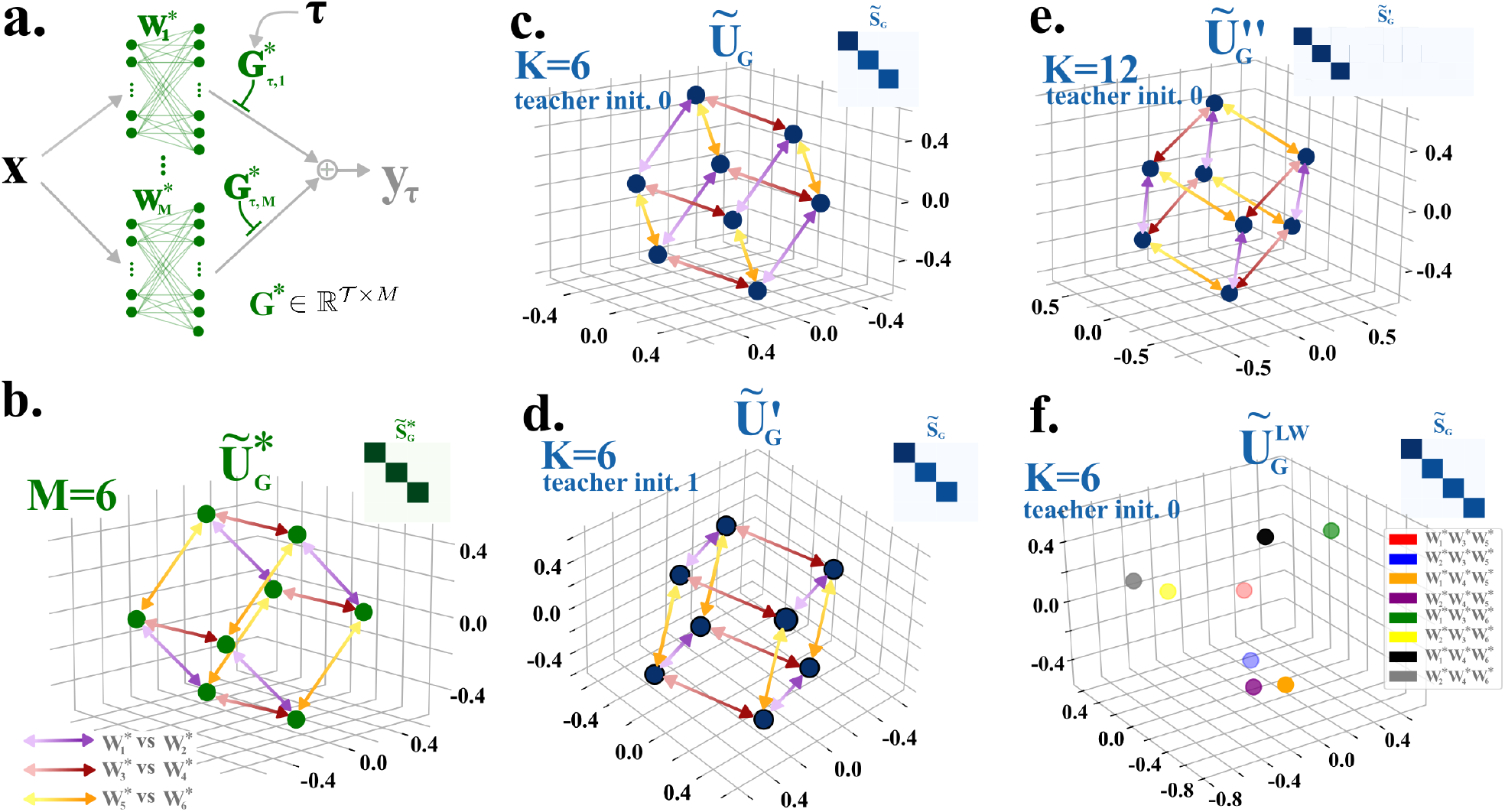
Networks Represent Compositional Structure in Gating Weights. **a)** Illustration of the teacher network used to generate training data. *τ* is an index which selects the row of **G**^∗^ that then gates the outputs of the set of teacher modules 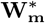. **b)** Visualization of the left singular vector matrix of 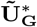. For each task *τ*, we plot 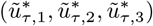, where *τ* corresponds to the rows of 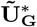. Since, 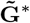 is rank 3 only the first three components of each row contribute to generating the overall values for each task. **c-f**) Same as **b**. but for the student’s left singular vectors, **U**_**G**_. **c** and **d**: *K* = 6, student initialized with small weights, and two different random initializations for the teacher weights, 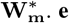: K=12, and student initialized with small weights. **f**: K=6, and student initialized with large weights. Since there are no consistent axes, we label each task individually.

The key analogy with the multitasking RNN is that the target of each task, **y**_*τ*_, is produced by combining outputs from different modules, 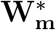, which, importantly, are reused across the task set, just as Mod1, Mod2, Strong, Weak, Resp1 and Resp2 are reused by the multitasking RNN. With a massive, but illustrative, abuse of notation, we can make this explicit by writing the outputs of the Mod1-Strong-Resp1 task as

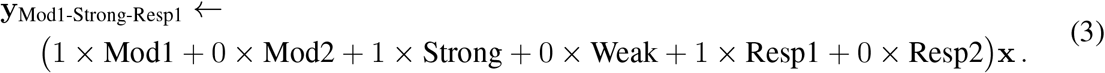

This captures the notion that the network combines the Mod1, Strong and Resp1 modules to perform the task. For the analogous task in our simplified setting, we take *M* = 6 in Eq. (2) and write

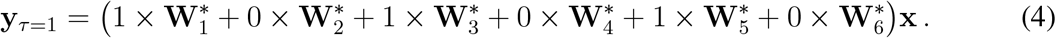

To draw the full analogy with the multitasking RNN, we let *T*= 8. The full task set structure is given by

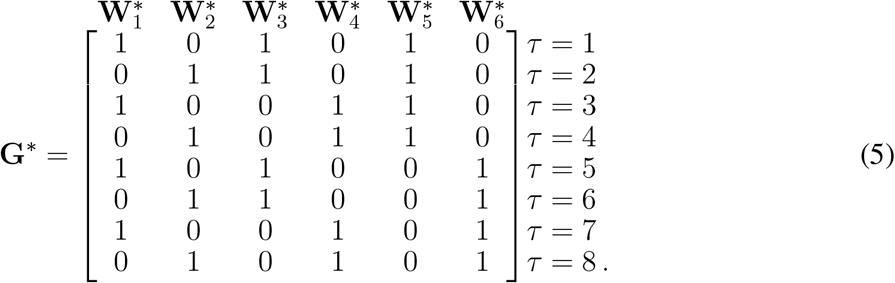

As indicated by the labels above and to the right of **G**^∗^, rows represent tasks, columns represent modules, and binary values indicate the inclusion or exclusion of a module in a given task. In this context *τ* is simply an integer that specifies which row of **G**^∗^ is active.

Like the example decision-making task set above, tasks in our linear setting are defined by 3 binary axes (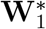 vs. 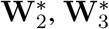 vs. 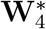, and 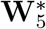 vs. 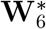). The resulting eight tasks are thus formed by three binary choices. Since **G**^∗^ is made up of zeros and ones, the binary choices are clear. In any row the columns corresponding to, for example, 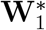 and 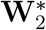, are always *either* 0 or 1 for a given task, never *both* 0 or *both* 1. Conceptually, then, each of these tasks lie on the vertices of a cube.

We are interested in determining the structure of learned weights in a network that has been trained on tasks produced by **G**^∗^. To do this we train a student network on data produced by the teacher network defined in Eq. 2. The student has a very similar architecture,

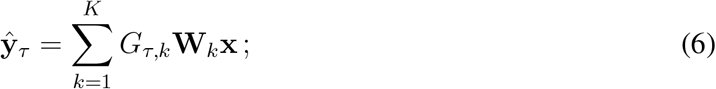

the only potential difference between this and the teacher architecture is that *K* may be greater than *M* (*K* ≥ *M*); i.e. the student is allowed to use more modules than were used by the teacher to produce the data. We train **G** and **W**_**k**_ to minimize mean squared error using gradient descent.

As we will see, the trained student networks will share this cube structure after learning a task set with this compositional structure, but the resulting weights are typically not binary. Hence, it will be useful at this point to make a precise mathematical characterization of the structure in **G**^∗^, which will in turn allow us to equate aspects of **G**^∗^ to weights of the student network at the end of training. In particular, let us consider the mean centered **G**^∗^, denoted 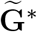 (this allows us to ignore the overall centroid of **G**^∗^ and focus on the internal relationship among tasks). The mean-centered contribution of each teacher module for a task *τ* is given by the row 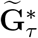. Now consider the singular value decomposition of 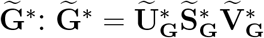. We can write each row 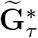 as

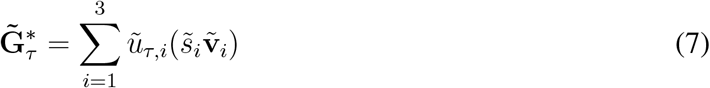

Note that in this case 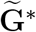 is rank 3, and there are only 3 non-zero singular values to sum over. Values *ũ*_*τ,i*_ are the first 3 components of the rows of 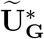. All of the structure of this task set is encoded in this formula. There are three axes 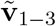 needed to describe the task set, and these correspond to the three binary teacher gatings (e.g. choosing either 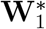 or 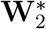, choosing either 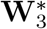 or 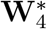, etc.). The components **ũ**_*τ*_ = (*ũ*_*τ*,1_, *ũ*_*τ*,2_, *ũ*_*τ*,3_) encode a given task’s position along these axes, capturing how each task is situated with respect to these binary gating choices. We plot **ũ**_*τ*_ for each task in Fig. 2b. Again, we find that each **ũ**_*τ*_ is a vertex of a cube. It’s also worth noting that for regular geometric structure like the one plotted in Fig. 2b to emerge, it is necessary that **G**^∗^ be low rank. A full rank **G**^∗^ implies that all directions are independent, which means no reuse and no shared axes (although in cases where *K > M* learned student **G** will be low rank even if **G**^∗^ is full rank; full rank **G**^∗^ here means that individual tasks may be distributed across multiple student modules, but those input-output maps will not be reused across tasks).

We can now outline how the compositional structure in the **G**^∗^ emerges in **G** after training. Full details of this analysis can be found in the Mathematical Supplement. For our purposes the key results are given by the following statements. Assuming the student network is initialized with small weights, then

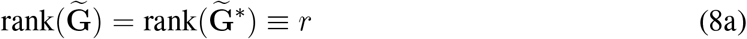

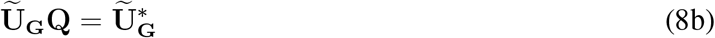

where 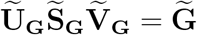 is the singular value decompositions of 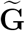 and **Q** is an *r×r* rotation/reflection matrix that acts on the rows of 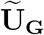 (i.e. the set of **ũ**_*τ*_). Intuitively, these equations simply state that the important structure we have already visualized in the data generating teacher is guaranteed to emerge in the student. First, Eq. 8a states that learned 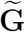 will have the same dimension as 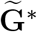. So if 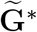 is low-dimensional, the learned 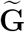 will also be low-dimensional. In compositional settings, a high degree of module reuse across the task set means this will often be the case. Second, Eq. 8b states that the geometry which captures how task components are reused across a task set defined by 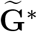 will be present in 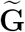 after learning. In our case, the task set is defined by three binary axes, which form a cube. This cube is explicitly encoded in the rows of the left singular matrix 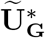. Equation 8b guarantees that 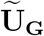 will form some rotated and reflected version of this same cube. Hence, the structure of the task is encoded in the geometry of the learned weights.

To illustrate these principles in simulation we trained four different students and visualized the set of **ũ**_*τ*_ (Fig. 2c-f). Again, these effectively serve as the coordinates of each task, and together reveal the geometry of task representations across the task set. The students shown in Fig. 2c-e have been initialized with small weights in accordance with our theory, while the student in Fig. 2f has been initialized with large weights. In Figs. 2c and 2d the student has *K* = 6, but for each model the weights, 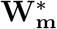, used to generate the training data were drawn using different random seeds, so each student learned a very different input-output mapping. Despite this, because the compositional structure of the task set is the same, the rows of the truncated left singular vector matrix still form a cube, as predicted by Eq. 8b. In Fig. 2e the student has *K* = 12 modules and was again initialized with small weights. Despite having twice as many modules as in panels c and d, a cube structure emerges. Again, this is predicted by Eq. 8b, which only requires there to be a sufficient number of modules in the student. These visualizations emphasize that the representations of module reuse encoded in 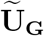 are independent of how individual maps are implemented (Fig. 2e). However, this structure is destroyed if the student is initialized with large weights, as in Fig. 2f, despite achieve the same performance on the task.

These plots show how the structure of module reuse across the task set is captured in 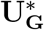. The crucial implication of Eqs. 8a and 8b is that when the student is faced with a set of compositionality related tasks, the learned **G** will be low-dimensional and will factorize tasks along dimensions that encode how modules can be reused. Here we chose to illustrate how structure is encoded in weights using a task with three binary axes because the resulting geometry – a cube – is intuitive given the organization of the task set. We emphasize, however, that Eqs. (8a) and (8b) give a general description of the structure of weights after training for any 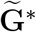 (given small weight initialization in the student). In other words, these equations provide a general formula that determines the geometry of task representations for any task set.

For the bi-linear network, all information about the current task is encoded in the rows of the matrix **G**. In fact, one can conceptualize these vectors as task representations analogous to task embedding vectors used in multitasking RNNs. In this way the set of weights **W**_emb_ in RNNs plays the same role as **G** in bi-linear networks. Based on this observation, we conjecture that **W**_emb_ will also contain the compositional structure of the task set after training. Quantitatively, that means the following. Suppose the structure of the task set is encoded in a matrix **G**^∗^ with the same dimensions as above. Let 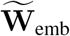 be the mean centered set of embedding weights and 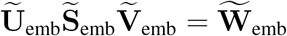 its singular value decomposition. Then

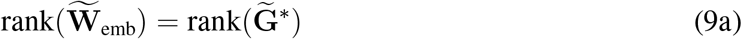

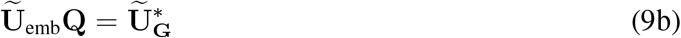

As above, these equations simply state that 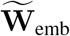 is low-rank, and the geometry of the left singular vectors will be inherited from **G**^∗^, which organizes the task set (i.e., in our worked example it will be a rotation and reflected version of a cube). In the next section we test Eqs. 9a and 9b empirically in nonlinear RNNs. If those equations are good predictors of the structure of learned task embedding weights in these more complicated models, then we’ll have a principled expression for what geometry will emerge after training. This will allow us in later sections to build algorithms for rapid learning from a solid foundation.

### Networks learn task set structure across classes and architectures

To test the conjecture summarized in Eq. (9), we trained RNNs on a variety of task classes in addition to the decision-making tasks outlined above. First, we trained models to output a sequence of cosine curves with different periods, which we refer to as the wave task (Fig. 3a). Previous work training non-human primates to produce sequences of periodic curves (among other shapes) showed evidence of reuse across similar shapes [42]. In this case, each individual cosine curve plays the role of a module, and individual tasks are built by concatenating these curves. Second, we trained networks on a Go task class (Fig. 3b). Like the decision-making task class, the Go task class has multiple input and response modalities. A correct response requires attending to the proper modality, responding either in the same direction as the stimulus (Go trial) or the opposite direction (AntiGo trial), and responding in the proper response modality. Finally, we trained networks on a ContinuousGo task class, which tests settings where tasks are made up of continuous interpolations of modules (Fig. 3c). These tasks have two input modalities and a single output modality. Participants must interpolate between the stimuli presented in Mod 1 and Mod 2, and respond with some interpolation between the resulting interpolation input angle and its opposite. For the Wave, Go, ContinuousGo task classes we use the architecture outlined in Fig. 1b.

**Figure 3:**
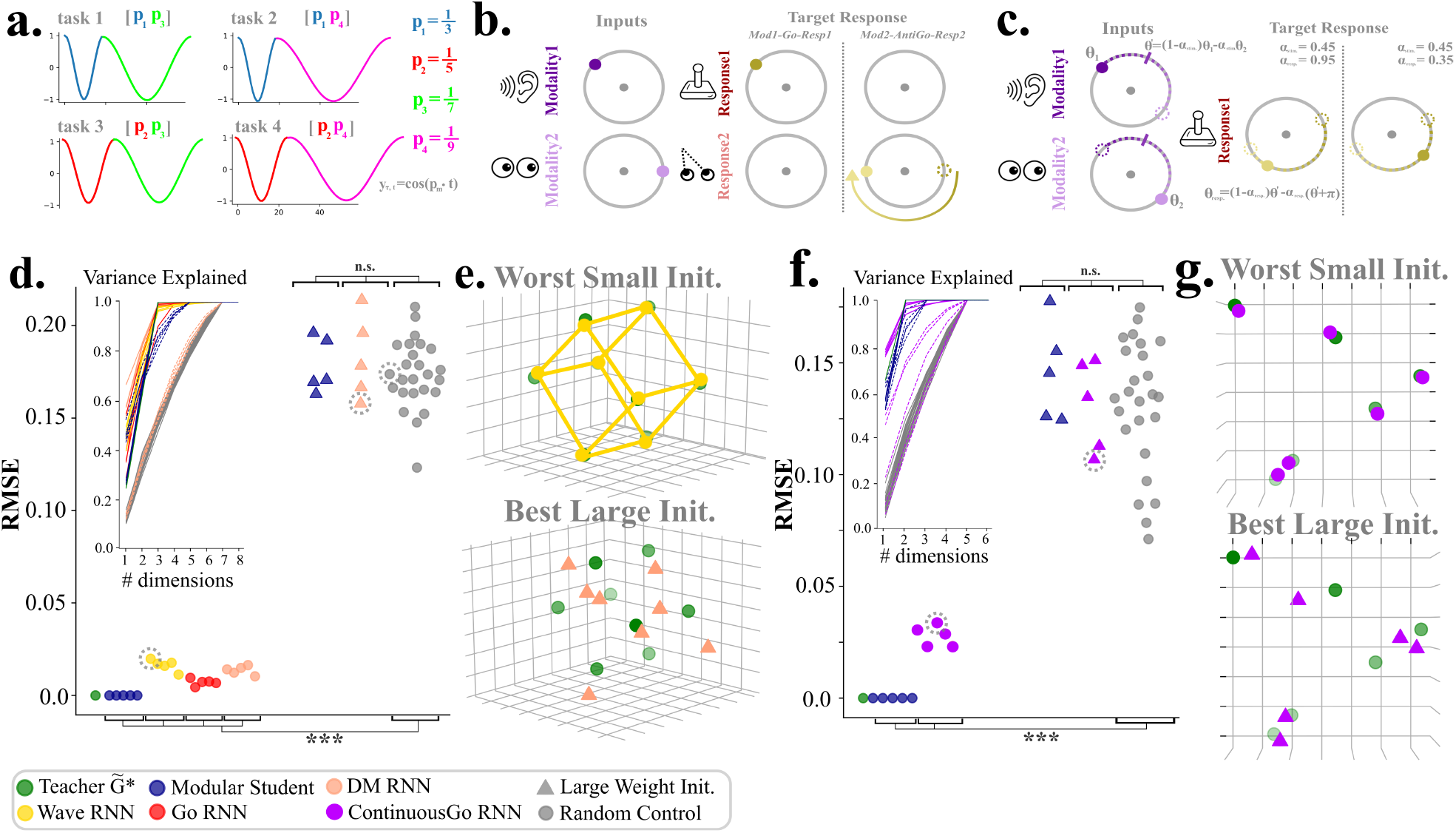
Learning task set structure across task classes. **a)** Illustration of four tasks from the Wave task class. Targets for individual tasks are built by concatenating cosine curves with different periods. **b)** Illustration of four tasks from the Go task class. Input and output occur in different stimulus and response modalities like in the DM task set. Otherwise, targets require a response to either the same (Go trial) or the opposite (AntiGo trial) of direction presented in the relevant stimulus modality. Example Mod1-Go-Resp1 and Mod2-AntiGo-Resp2 trials shown. **c)** Illustration of two example trials from the ContinuousGo task class. Each task is defined by two parameters. First, *α*_stim_ defines the degree of interpolation between the stimuli presented in the two sensory modalities. Second, *α*_resp_ determines a the degree of interpolation between the interpolated input and its opposite, which defines the target output. **d)** Correspondence between teacher 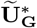 and learned **Ũ**_**G**_ and 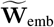 across task classes. For each we find the best rotation between learned weights and the teacher, and measure the RMSE between the resulting values. Dots show models trained with small weight initializations and triangles show models with large weight initializations (***: *p <* 1*e* − 4, corrected Mann-Whitney Test). Different colors indicate the training task class, and each point represents a model initialized with a different random seed. Variance explained inset shows the cumulative variance explained by each singular value of the relevant weight matrices. Dotted lines indicate models with large weight initializations **e)** Visualization of **Ũ**_emb_ rotated onto the teacher 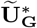. Dotted circles in **d**. indicate which models representations are being plotted. In both top and bottom panel the ground truth task representations from **G**^∗^ are plotted in green. Top panel shows the model with small weight initialization that has the worst RMSE. In this case the worst small initialization model is trained on the wave task, and the cube is colored accordingly. Bottom panel shows model with large weight initialization with best RMSE, here colored for the DM task class. **f)**-**g)** Same as **d**-**g** but for tasks with continuous interpolation of modules.

To begin we formulate task sets for the Wave and Go task classes that are organized by three binary axes, in analogy to the examples outlined above (see Methods for details). We trained models to convergence on all 8 tasks for the Wave, Go, decision-making task classes. Once the models are trained, we determine whether 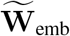 has approximately the same rank as 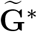, in accordance with Eq. 9a. To do so we measure the proportion of variance explained by each of the eigenvalues of 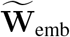. Further, recall that we that 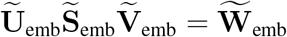 and 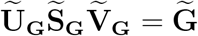. To determine whether the rows of **Ũ**_emb_ lie on the vertices of a cube in accordance with Eq. (9b), we find the best rotation between 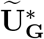 and **Ũ**_emb_ and take the root mean squared error (RMSE) between the resulting set of points (see Methods, Empirical Tests of Compositional Structure). For controls we take random matrices with the same dimensions as **W**_emb_ and perform the same analyses.

The results are shown in Fig. 3d. To ground our empirical results, we first plot each of these measures for student-teacher networks where analytical results guarantee low-rank structure and zero-distance rotation measure. This results in zero RMSE and only three important dimensions (Fig. 3d., modular student). Next we examined these measures in the embedding weights of RNNs. Strikingly, these also revealed that when initialized with small weights, **Ũ**_emb_ is also approximately a rotation away from the 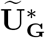, in line with the conjecture given in Eq. 9b. This correspondence is destroyed when the RNN is initialized with large weights (triangles). The inset shows the cumulative variance explained by successive singular values. For all models initialized with small weights only three eigenvalues of 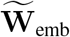 are required to explain over 90% of the variance, implying that 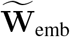 is effectively rank 3, in accordance with the conjecture in Eq. 9a. A visualizations of **Ũ**_emb_ rotated onto teacher 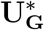 is shown in Fig. 3e. The top panel shows the optimal rotation for the model with the worst RMSE and small weight initialization, and the bottom panel shows the optimal rotation for the model with the best RMSE and large weight initialization.

Analogous results are shown in Fig. 3f for the ContinuousGo task set. In this case each task is defined by two variables: *α*_stim_ and *α*_resp_ (see Fig. 3c). Unlike the previous tasks, this set is not structured along three binary axes; instead, it is parameterized by two continuous variables. Intuitively, task representations here should simply reside at coordinates on the 2-D plane that correspond to parameter values. With our conjecture we can validate this exactly. Indeed, we find that when the RNN is initialized with small weights, Eqs. (9a) and (9b) predict the structure in learned weights: **Ũ**_emb_ is approximately a rotation away from 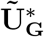, and the ranks of the two matrices are approximately the same. This is illustrated in Fig. 3g, which shows the worst rotation for small weight initialization and the best for large weight initialization.

Finally, we tested three other task classes, each with different compositional structure. Across these configurations we found that Eqs. (9a) and (9b) provided a good approximation to the structure of weights after learning (see Supplementary Fig. 1). Crucially, in one of the tested configurations we held out a task from our three binary axis task set. In this case the resulting geometry is a cube with one vertex missing.

The results in this section provide strong empirical evidence that Eqs. 9a-9b are a good approximation of the structure in **W**_emb_ after learning. This gives us a general formula that predicts the geometry of neural representations given the organization of the task set. This is a step towards explaining similar geometries in artificial neural network activations and in neural recording of animals engaged in tasks with overlapping demands [25, 33, 34]. For the purposes of rapidly learning new tasks, these results are crucial because they show that even in cases when tasks are held out of training, the network will extract the key structural organization of the task set (Supplementary Fig. 1e.-f.). Now that we have a mathematical description that approximately characterizes this learned structure even in complicated tasks and architectures, we are in a position to build algorithms which networks can reliably use to constrain, and therefore speed up, learning of new tasks.

### Rapid Learning Using Compositional Structure and Reinforcement

The embedding weights, 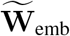, encode the compositional structure of the training set (given small weight initialization). In particular, we found empirically that 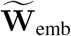 will be low-dimensional, and the structure of the task set will be encoded in the geometry of **Ũ**_emb_. The key implication of these facts, and the core thesis of this work, is that learning for a novel but compositionally related task can proceed in a subspace of activity which is significantly smaller than the full task embedding space. In this section we show that learning in this constrained space will result in very rapid acquisition of new tasks.

Let us return to our DM task set with three binary components. Suppose we have a model trained on seven of the tasks with the Mod1-Strong-Resp1 task held out, as depicted in Fig. 1. Now, we want to learn the held out task (for brevity and generality we’ll refer to the held out task as *τ*_*h*.*o*._). In the setting we consider, no task cue information is provided to the model, but it receives a reward for a correct response. Importantly, the system also does not know that the target task is novel and so has no reason to exclude training tasks from exploration. The most unconstrained method of learning that still takes advantage of knowledge acquired during training is the modular learning, discussed briefly in **Modularity and task set structure in multitasking recurrent networks as a path to modeling rapid learning**. This method involves freezing weights in the RNN and updating only the activity of the task embedding (denoted 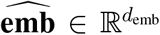). This is illustrated in Fig. 4a, which shows a cartoon of a hypothetical learning trajectory. The trajectory freely explores all *d*_emb_ dimensions of the task embedding activity space.

**Figure 4:**
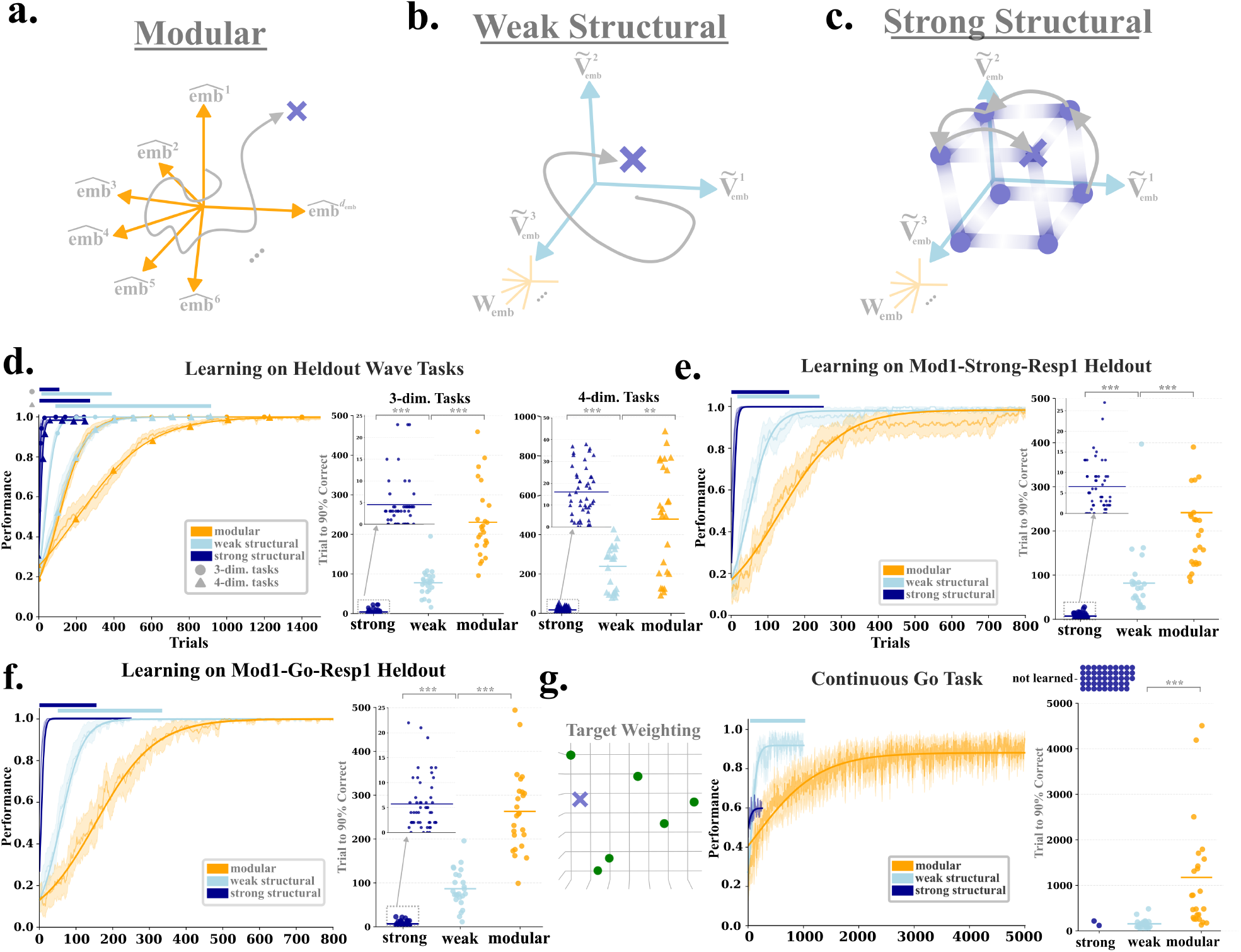
Learning from reinforcement using the structure of compositional tasks. **a)** Illustration of modular learning. A reward signal is used to update a vector of activity in the full task embedding space (orange axes). Hypothetical learning trajectories are plotted using gray curves, where the blue x marks a region of high reward for the target task. **b)** Illustration of weak structural compositional learning. Here we update activity in a low-dimensional control signal **c** (gray trace) which determines the location of the task embedding within the relevant basis 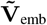 (blue arrows) determined by learned **W**_emb_ (see Methods for details). **c)** Strong structural composition takes advantage of the geometric regularities of task embeddings that emerge in task sets which involve module inclusion or exclusion. Here exploration during learning proceeds according to a strong prior (shaded blue regions) which preferentially explores the vertices of the underlying parallelogram (in this case the cube). **d)** Learning curves for a held out task on the Wave class of tasks for both the 3 binary axis cube structure (circles) and an analogous 4 binary axis hypercube structure (triangles). Shaded regions show +/− s.e.m. Modular learning is shown in orange, weak structural composition (w.s.) in light blue and strong structural (s.s.) composition in dark blue. Dark blue bars above learning curves indicate where s.s. is greater than w.s. and modular learning by a statistically significant margin, as assessed by cluster-based permutation test (see Methods, Compositional Learning from Reinforcement). Light blue bars indicate where w.s. is greater than modular according to the same test. On the right, scatter plots show the number of trials required by each learning run to reach 90% performance threshold for each learning method (3-dim. tasks: w.s.>s.s U=1247.0, p=6.61*e*^−12^; modular > w.s. U=611.0, p=2.21*e*^−8^; modular > s.s. U=1250.0, p=4.83*e*^−12^. 4-dim. tasks: w.s. > s.s. U=1250.0, p=6.61*e*^−12^; modular > w.s. U=477.0, p=4.38*e*^−3^; modular > s.s. U=1250.0, p=6.61*e*^−12^, corrected Mann-Whitney Test). **e)** Same as **d)** except for DM task with three binary axes; Mod1-Strong-Resp1 held out (threshold stats: w.s. > s.s. U=1250.0, p=6.23*e*^−12^; modular > w.s. U=565.5, p=2.87*e*^−6^; modular > s.s. U=1250.0, p=6.25*e*^−12^). **f)** Same as **d)** except for Go task class; Mod1-Go-Resp1 held out (threshold stats: w.s. > s.s. U=1241.5, p=1.17*e*^−11^; modular > w.s. U=612.0, p=1.96*e*^−8^; modular > s.s. U=1250.0, p=5.94*e*^−12^). **g)** Same as **d)** except for the ContinuousGo task with the same structure as shown in Fig. 3e. The position of the target task relative to the embeddings is shown in the left most inset. Here, strong structural composition fails to learn the task in the majority of runs, which are plotted in the scatter as ‘not learned’ (threshold stats: mod. > w.s. U=576.5, p=6.61*e*^−12^).

Past work using similar sets of tasks has shown that this method can effectively recruit the proper recombination of modules in the RNN to perform novel tasks [21]. However, the empirical results in the previous section show that across different task types and task set organizations there is an enormous amount of information about the compositional structure of the task set in the learned weights **W**_emb_ which modular learning ignores. For instance, we showed that the dimension of the task set structure determines the rank *r* of 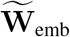. Hence, to perform a related task within this structure, we can forgo exploring the entire embedding activity space and constrain learning to the subspace defined by the basis of the mean centered embedding weights 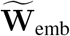 (note we can trivially add back the mean at any point). This method ensures that the dimensionality of the learning problem is determined by the complexity of the task set, rather than arbitrary choices about the architecture of the model. In this setting, we simply learn a vector of activity **c** ∈ ℝ^*r*^ which determines the location of the embedding in the relevant basis. We call this weak structural composition because it takes advantage of the low-dimensionality of the embedding space, but does not leverage any geometric regularities of the task embeddings themselves. A cartoon illustration of a hypothetical learning trajectory is shown in Fig. 4b. In this cartoon, learning is constrained to the light blue axes, which form the relevant subspace of **W**_emb_.

We can learn even more efficiently by taking advantage of geometric regularities in the task embeddings. In particular, if **G**^∗^ is binary (i.e., modules are either included or excluded from a task), then the resulting task embeddings lie on a subset of the vertices of an *r*-dimensional parallelogram (see Mathematical Supplement). The highly regular structure of these embeddings means that, given a sufficient number of training tasks, we can infer the position of held out tasks embeddings by filling out the vertices of the underlying parallelogram. We can then formulate a strong prior over exploration which preferentially explores this set of vertices. A hypothetical learning trajectory is shown in Fig. 4c. Blue shading in this cartoon represents the policy prior formulated from the geometric regularities of the embedding space. Vertices of the cube are more likely to be explored, and this space includes the correct embedding (shown with an *×*) for the held out task.

For both modular and weak structural composition, learning proceeds using the REINFORCE algorithm with 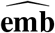 and **c** parameterized by normal distributions. Note that the goal of learning in all cases is to find a continuous vector of activity that maximizes reward (as opposed to weights of a deep neural network). For strong structural learning we map the original embedding space onto the *r*-dimensional hypercube aligned to the unit axes so that each task resides on a vertex of the hypercube. In this case, our control vector is parameterized by a multi-dimensional Beta distribution, with concentration parameters *α* and *β*. Reward feedback is used to update these concentration parameters (see Methods, Compositional Learning from Reinforcement). Crucially, at the beginning of learning we set these *α* and *β* to be small. This induces a prior on **c** with high density around 0 and 1 along each dimension, which then preferentially samples embeddings at the vertices of this hypercube. For all methods we update parameters after reward experienced on a single trial, matching the feedback received by animals in many experimental settings (see Methods, Compositional Learning from Reinforcement).

Results are plotted in Figs. 4c-f. Figure 4c. shows learning curves for the held out wave task with with 3 binary axes (like in Fig. 3a., here denoted 3-dim tasks) and 4 binary axes (denoted 4-dim task). On the right of the learning curves we plot the number of trials required to reach 90% correct for each learning run. As expected, modular composition exhibits the slowest learning because it ignores the low-dimensional structure of task embeddings. By contrast, constraining learning to the relevant subspace of task embedding activity in weak structural composition results in learning that is approximately twice as fast as modular composition. Finally, the prior used in strong structural composition successfully guides learning such that models reach 90% in 5 trials on average for 3-dim tasks and 16 on average for 4-dim tasks. This represents a full order of magnitude increase in required learning time for acquiring the novel tasks. Figures 4d and e show the same learning information for the DM and Go tasks respectively, and show roughly the same relative increase in the speed of learning across methods.

The speed of learning in strong structural composition arises from the strong assumptions made about the training set; namely, that each task is purely a matter of module inclusion or exclusion. If this is indeed the case, the full embedding space will form the vertices of a *r*-dimensional parallelogram which can be preferentially explored, and learning will occur in a handful of trials. We note that this assumption holds for a large swath of experiments designed to probe compositional abilities, making strong structural composition a widely applicable model [29, 31, 33]. However, the assumptions required for strong structural composition to succeed do not hold in any arbitrary setting. In particular, when tasks are continuous interpolations of module computations, embeddings will not form a parallelogram, and strong structural composition should fail to learn the task. An example is shown in Fig. 4g, where we learn an unseen task in the ContinuousGo class. This task lies in the plane defined by the embeddings for the training tasks in Fig. 3g, top panel (target task marked by the blue *×* in the inset of Fig. 4g. In this case, strong structural composition fails to learn the novel task in 90% of learning runs. The only structural constraint that is guaranteed in this setting is that task embeddings overall still reside in the 2-D plane. This means that weak structural composition can still usefully guide learning. This method reaches 90% performance in 151 trials on average, still highly efficient compared to modular learning which is an order of magnitude slower (1,137 trials to threshold).

To demonstrate the core principles of constraining learning using compositional structure, we examined task sets with a single tasks held out from a multitasking RNN. We perform a number of follow up analyses that are include in Supplementary Information. We examine the differing reward landscapes in models trained on Go vs. ContinuousGo task sets in Supplementary Fig. 2. In Supplementary Fig. 3, we show that models with multiple tasks held out of training can still develop the representational organization required to support this rapid of learning, and outline the specific conditions required for these representations to emerge. We show that updates to 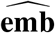 in weak structure learning are much more likely to follow the true gradient of this vector than in modular learning (Supplementary Fig. 4). We now turn to a more detailed examination of learning via the geometric regularities in strong structural composition.

### Strong structural composition approximates discrete hypothesis testing over a continuous activity space

Learning in the case of strong structural composition involves placing a strong prior over the vertices of the underlying parallelogram that scaffolds task embeddings. The strength of the prior means that during learning, exploration effectively takes place over a discrete set of points. In this section we demonstrate that strong structural learning shows many of the characteristics of learning via hypothesis testing.

A hallmark of learning via hypothesis testing is discontinuous learning curves, where individual participants shows a sharp and sudden jump in performance rather than a steady increase. Past behavioral work has observed these dynamics in experiments, but note that these characteristics are often obscured by averaging learning curves over a population. This results in a much smoother curve overall [5]. We see exactly these dynamics when examining learning curves for individual runs in strong structural learning (Fig. 5a). The sudden jumps in performance shown in this figure occur because the control vector **c** jumps between the vertices of the cube, rather than moving smoothly through a low-dimensional space. To further corroborate this we plot the euclidean distance between **c** and the target control vector **c**∗ for each learning run. On unrewarded trials **c** moves between distances of either 1, 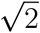, or 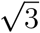, corresponding to an adjacent vertex, a vertex diagonal on the same face, and an antipodal vertex. This amounts to hypothesis testing across the set of tasks, where on each trial the system attempts a combination of the underlying modules, collects reward or not, and based on this outcome either proposes an entirely new combination on the very next trial or maintains the rewarded combination from the previous trial (for a more complete characterization of learning dynamics after both rewarded and unrewarded trials see Supplementary Fig. 5).

**Figure 5:**
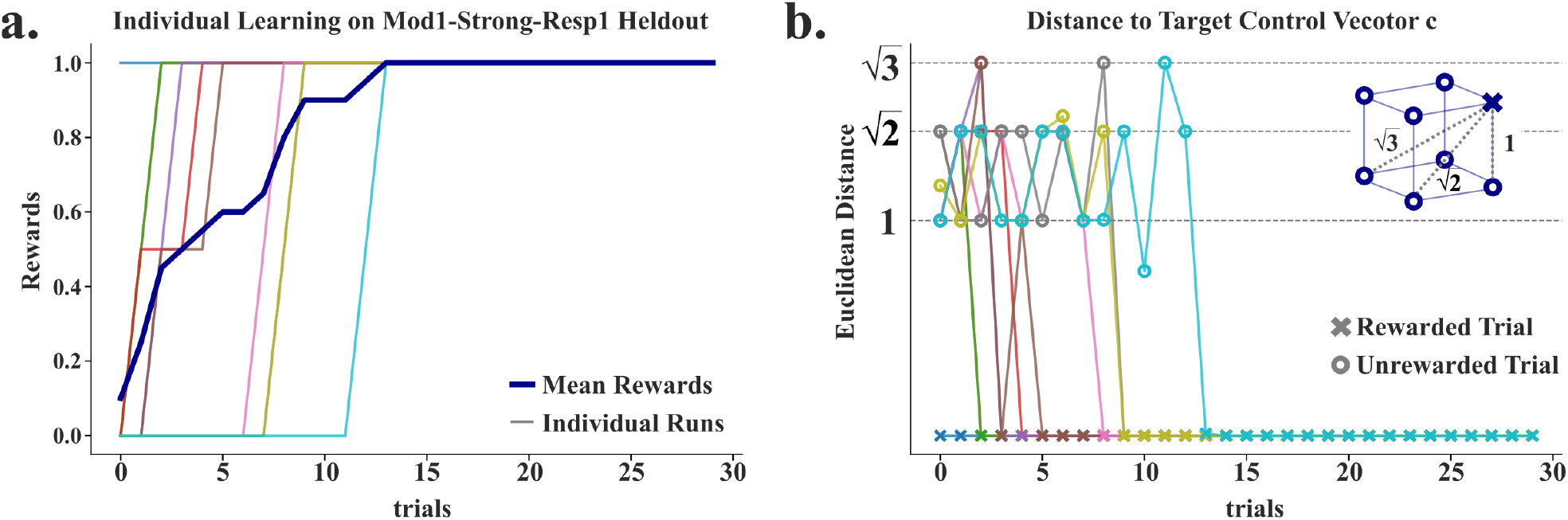
Strong structural composition implements hypothesis testing style learning. **a)** Learning curves from strong structural learning on Mod1-Strong-Resp1 task. Dark blue curve shows learning averaged across the population and colored curves show learning for 10 individual runs from a model seed (plotted as a moving average with window size 2). **b)** Distance between the control vector **c** at each point in learning and target vector **c**∗ for each individual learning run. Distances cluster around 1, 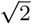, and 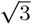 which correspond to adjacent vertices, vertices diagonal on the same face, and antipodal vertices, as shown in the inset. Open circles indicate unrewarded trials and x’s indicate rewarded trials.

### Capturing neural signatures of compositional task discovery

In the previous sections we showed how weights in neural networks encoded the structure of module reuse across a task set, and how leveraging that structure allows for very rapid acquisition of novel tasks. At a basic level our models describe how neural activity should evolve if agents are using the compositional structure of the task set to guide learning. However, the speed of learning exhibited by structural composition opens the possibility of fitting these neural models to the behavior of animals engaged in compositional tasks, where adaptation to task demands can be very rapid [29]. We can then compare the evolution of activity in our models to data. As we will see, constraining learning with compositional structure makes specific predictions about how modules are re-engaged as learning occurs, which we find is a good match to trends observed in data. In this section we first describe the experiment we intend to model, and outline two alternative models which we fit to behavior alongside structural composition. From there, we show how learning via structural composition is the only model which can capture the important features of neural evolution during the acquisition of task demands.

In a recent experiment testing animals’ compositional abilities, Tafazoli et. al. [29] trained two monkeys on a set of decision making tasks defined by 1) the relevant feature of the stimulus and 2) the relevant axis of responses. On the feature dimension, the monkeys had to perform a 2-way categorization based on either the shape of the stimulus (more ‘Tee’-like or more ‘Bunny’-like) or the color of the stimulus (more red or more green). The animals then had to report their answer to the decision-making task using the proper response axis. For response axis 1 trials, animals had to answer using top-left and bottom-right screen touches; for response axis 2 trials, they had to respond with top-right and bottom-left screen touches. Animals were pre-trained on the shape-axis1 (S1), color-axis1 (C1), and color-axis2 tasks (C2) only (Fig. 6a). The decision boundaries for both shape and color features are shown in Fig. 6b. After pre-training, the animals achieved high performance in each task individually. The authors then recorded neural activity across multiple brain regions while the animals transitioned through a sequence of blocks, where trials in each block are from a single underlying task type (Fig. 6b, bottom). Block transitions were explicitly cued, but the cues provided no information about the identity of the upcoming task. In this setting, the main task of the animals is to infer the current task demands using reward feedback alone. This paradigm allowed researchers to track neural dynamics during “task discovery”; i.e., inference of current task identity. Though contextual inference has been extensively studied in previous work [30, 43–45], this experiment is the first to record from non-human primates in a task set where contexts are made up of distinct compositional components (stimulus feature and response axis). Although in modeling this experiment we shift focus from learning novel tasks to discovering previously learned tasks, the design allows us to test our theory of compositional learning (as opposed to many other contextual inference tasks which lack compositional structure). The basic proposal of our model is that when the system is attempting to configure modules to find a compositional solution to a task, the process of selecting modules should be confined to low-dimensional space defined by the structure that emerges in embedding weights. In the context of this experiment, our theory is that these dynamics will govern how animals discover the underlying task in any given block, even if the animal has previous experience with the task.

**Figure 6:**
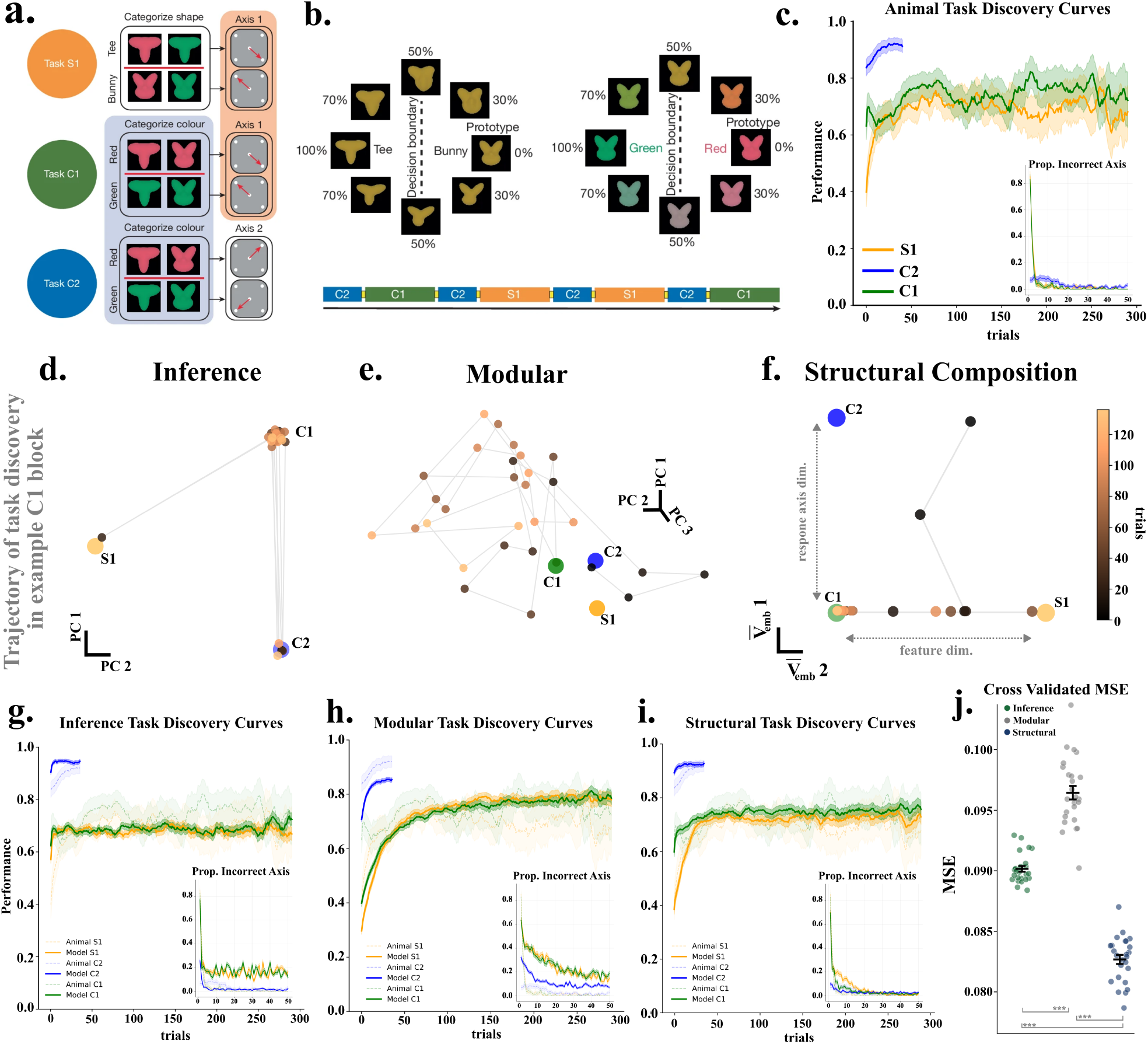
Structural composition captures key neural features of task discovery. **a**. Space of stimuli and decision boundary in the shape decision-making task (top) and color decision making task (bottom). For clarity, color is kept constant in the shape examples and vice versa in the color examples. In actual trials both shape and color vary. Adapted from Tafazoli et al. **b**. Examples of the three kinds of tasks included in the experiment. In S1 the participant must categorize the stimulus according to its shape and respond with the first axis (top-left, bottom-right). In C1 the participant must categorize the stimulus according to its color and respond with the first axis. In C2 the participant must categorize the stimulus according to its color and respond with the second axis (top-right, bottom-left). Below is an illustration of the sequence of blocks that is experienced by the animal in the experiment. Adapted from Tafazoli et al. **c**. Learning curves averaged over animals and blocks for different task types (smoothed over a sliding window of 10 trials). Inset shows the average proportion of trials where animals responded with the incorrect response axis as each block type progresses. **d**. Trajectory of task discovery in C1 block for fitted Inference Model. Large labeled points indicate the location of task embedding vectors established during pre-training for each task. Individual points represent the embedding vector every 5 trials for the first 150 trials of the block. For Inference model a slight jitter is used to visualize otherwise overlapping embeddings **e**. Same as **d**. on the same C1 block but for fitted Modular Model. **f**. Same as **d**. on the same C1 block but for fitted Structural Model. Here axes are unit aligned right singular vectors of 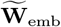 corresponding to feature dimension and response axis dimension of the task set. **g**. Fitted learning curves for the Inference Model (cross validated parameters, averaged across 5 pretrained RNN initializations and fits for 2 animals) plotted against animal learning curves. Inset shows the average proportion incorrect response axis as the block progresses. **h**. Same as **g**. but for Modular Model. **i**. Same as **g**. but for Structural Model. **j**. Cross Validated MSE between animals’ and models’ fitted task discovery trajectories. Points represent 5 runs of the model for 5 RNN initializations, averaged over animals (***: p«0.05; corrected Mann-Whitney Test).

Experimental learning curves over both animals in each task type are shown in Fig. 6c. For our purposes, the key feature of behavior is that animals are initially very quick to learn the current task demands, achieving well above chance performance within the first tens of trials. This is followed by a more gradual increase in performance as the block progresses, specifically in S1 and C1 blocks. Part of what drives the initial speed in these learning dynamics is very fast discovery of the current response axis. The inset of Fig. 6c shows how often animals responded using the incorrect axis as the block progressed. The number of error drops dramatically within the first few trials. One important aspect of the experimental design is the sequence of blocks. In particular, a C2 block is always followed by an S1 or C1 block, and S1 and C1 blocks always transition to C2 blocks. There is therefore, in principle, enough information for animals to do very well on the first trial of C2 block, and that there is less uncertainty in these blocks because animals don’t have to adjudicate between shape and color as the relevant feature (this likely accounts for differences in ceiling performance between block types). Animals sometimes take advantage of block sequencing information. We include parameters in our models that capture these sequence effects though we don’t attempt to model sequence learning in a mechanistic way, and instead focus on the dynamics of task discovery within individual blocks (see Methods). For possible mechanisms capturing this block sequencing see [46].

Here we fit our reinforcement learning models to the task discovery trajectories of animals [5, 47]. We pre-train a multitasking RNN so that it is proficient on all tasks individually (mirroring animal pre-training) and use these networks to model animals’ task discovery trajectories. To model animal behavior we pre-train models on the same three tasks (S1, C2, C1; training on C1, C2, S1, and S2 yields a square geometry in task embedding vectors where axes are defined by the feature dimension and the response axis dimension, Supplementary Fig. 1). Following the kinds of learning discussed in previous sections, we assume that the RNN recurrent weights which implement modular computations are frozen, and instead we model task discovery as primarily a result of changes in embedding activity which serves as external input to the RNN itself.

Given that the animals have observed all tasks during pre-training, the most straightforward way to model task discovery is as an inference process, where the animal uses reward to update its belief that a given task is the true underlying task in that block. At each trial a task embedding is sampled from this distribution (we call this the Inference Model, for full details see Methods). An example embedding trajectory in a C1 block for a fitted Inference Model is shown in Fig. 6d. In this model, because the space of possible embeddings is very small (three points acquired during pre-training) learning can be very fast. However, it is fundamentally different than the compositional learning we have been discussing because it does not learn according to the task relevant dimensions that emerge in embedding weights, and instead simply updates a distribution over the known set of tasks.

Alternatively, animals might discover task demands by updating a the full high-dimensional task embedding vector. This is exactly the modular learning consider in previous sections. Figure 6e depicts a trajectory of embedding activity in the same block as Fig. 6d for a fitted Modular model. Here activity freely evolves in high-dimensional space, meaning some learning occurs along the relevant dimensions, but also can occur in orthogonal and irrelevant directions.

Finally, Fig. 6f shows the trajectory of learning in the same block for a model using structural composition (because in this case we don’t impose a strong prior on exploration but still parameterize the policy using a beta distribution, we drop the strong/weak distinction and simply refer to this model as structural composition; see Methods for details). In this case we plot the evolution of low-dimensional control vector **c** which gets mapped back into the full embedding space. Again, in this case **c** is 2-dimensional, with one dimension corresponding to the task feature (shape vs. color) and another corresponding to the response axis (axis1 vs. axis2). Because activity only evolves along the task relevant feature and response axis dimensions this model can accommodate fast acquisition of the proper response axis and a more gradual convergence on the color feature.

We use Bayesian optimization to minimize the mean squared error between animals and model task discovery trajectories (defined as the performance averaged over a moving window of 5 trials) across all blocks in the dataset (full details can be found in Methods). The resulting per rule task discovery curves for each model are shown in Fig. 6h-j. The inference model learns very rapidly but stays flat through later trials in the block. Modular models learn more gradually in later block trials, but fail to capture the fast learning exhibited in early trials. Structural composition, by contrast, captures both the initial rapid rise in performance and the gradual increase in C1 and S1 rules as the block progresses. At a high level this for the same reasons illustrated in Fig. 6f.; structural learning can implement very fast response axis learning and more gradual feature axis learning in the same model because each task dimension is naturally represented separately, and learning takes place exclusively along these directions. Fig. 6j shows the cross validated MSE between model and animal task discovery curves across the entire dataset (points represent 5 runs from 5 RNN random initializations, model fits for each animal individually are shown in Supplementary Fig. 7).

**Figure 7:**
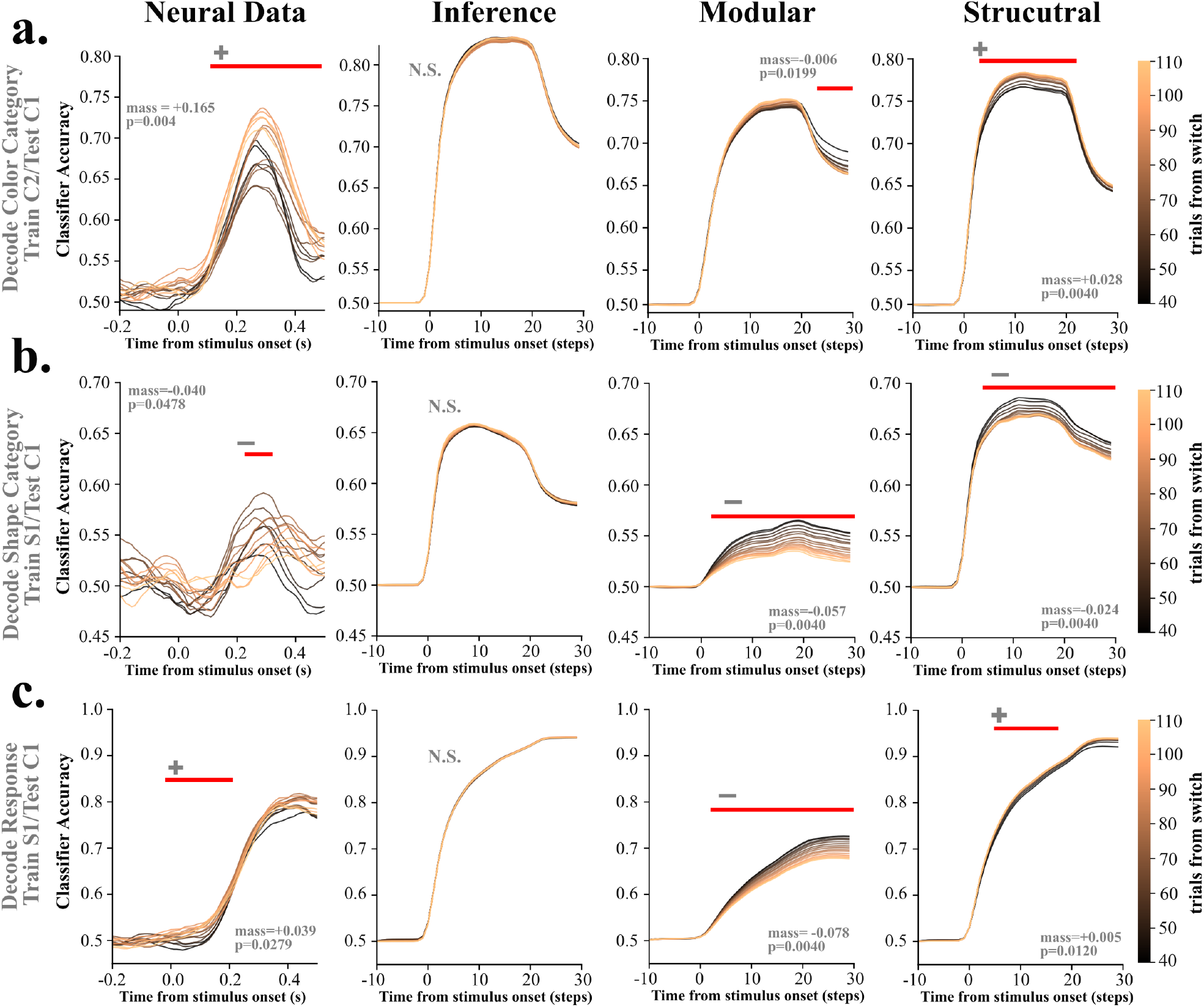
Structural composition captures key neural features of task discovery. **a**. Cross decoding performance on color feature. Decoders were trained to discriminate the color category of the stimulus from the activity in trials at the end of the C2 block. These decoders were then tested on sliding windows of 40 trials from C1 blocks. Red bars indicate a statistically significant trend in decoding performance as the C1 block progressed (see Methods, Cross Decoding Trend Test). The *±* over the bar indicates the direction of the trend. First column shows neural data, second shows the same analysis from fitted Inference model, the third for the Modular model, and the fourth column show cross decoding for the structural model. **b**. Same as **a**. but for a decoder trained to discriminate the shape category of the stimulus from the final trials of S1 blocks, then tested on shape discrimination on the initial trials of C1 blocks. **c**. Same as **a**. but for a decoder trained to discriminate the response direction from the final trials of S1 blocks, then tested on response direction on the initial trials of C1 blocks.

Structural composition allows us to fit the often rapid task discovery trajectories of animals. Because trials responses at each stage of task discovery are the result of neural computations in the RNN, we can probe the extent to which this behavior is the result of similar mechanisms as those observed in neural data. The most relevant analysis examining how representations evolve during task discovery in Tafazoli et al. tests how decoders trained on lateral prefrontal cortex (lPFC) activity generalize from the end of blocks in one task to the beginning of blocks in an alterative task. We replicate these analyses using decoders trained on RNN activations. First, we trained decoders to differentiate the color category using trials from the end of C2 blocks. We then evaluated these decoders on the first 110 trials of C1 blocks. Tracking the trend of cross decoding performance reveals how different computational subcomponents are re-engaged as task discovery progresses. Results are shown in Fig. 7a. In neural data, cross-decoding performance improves steadily as the animal advances through the block, indicating there is re-engagement of the relevant color-discrimination module. Next, we tested a decoder trained to differentiate shape category in S1 blocks on trials from C1 blocks (Fig. 7b). In this case accuracy in neural decoding shows a negative trend as the block progresses, reflecting animals disengagement of shape discrimination modules. Finally, we tested a decoder trained on response direction in an S1 block on trials from C1 block (Fig. 7c). Here, neural decoding shows a slight positive trend. Animals re-engage response axis 1 module, but do so very quickly, resulting in a smaller trend overall. Inference models fail to capture any of these trends. Although cross-decoding is high in this model, the speed with which inference models discover the C1 task means there is no statistically significant trend as the block progresses. Modular learning shows the proper trends for the shape decoding, and an insignificant increase in color decoding accuracy during the stimulus period (time steps 0-20) followed by a decreasing trend in the response period (time steps 20-30). Most notably the modular model shows a large negative trend for response axis cross decoding. This is likely the result of unconstrained learning. Because task embeddings are free to move in any dimension, it is possible for RNN activity to produce correct responses that nonetheless drift away from the format of response axis representations at the end of S1 block where the decoder was trained. The only model that correctly reproduces the trends observed in data across all three conditions is the structural model. Here constraining task discovery to occur in the relevant subspace means modules in the RNN are reliably reactivated as task embeddings move along the task relevant axes. This results in gradual re-engagement of color, disengagement of shape, and fast re-engagement of response axis modules.

Overall, these results show that constraining task discovery to occur along the dimensions discovered by embedding weights leads to both behavior and neural dynamics that are consistent with experiments. This is evidence toward the possibility that learning along dimensions defined by the structure of module reuse also occurs in animals engaged in compositional tasks.

## Discussion

The core claim of this work is that a compositional system which leverages the structure of module reuse will be able to acquire unseen tasks very rapidly. Our analysis of composition in bi-linear networks showed that this structure necessarily emerges in learned weights given small weight initialization. We then conjectured that more complicated non-linear recurrent systems behave in the same way, and validated this empirically over a variety of task types and compositional structures. Using this structure allowed us to constrain learning, resulting in very rapid task acquisition. Finally, we offer initial evidence that monkeys are using the structure of module usage to guide learning (or in this case task discovery). These results capture a long held intuition taken from classic cognitive science: that much of learning involves recombining pre-existing knowledge in novel ways, rather than starting from scratch at every new problem. The key contribution of this work is to show that under very mild conditions exactly this kind of learning is very easy to do in a system of neurons, precisely because they neurons will encode a simple representation of which recombinations are possible. These representations then can then efficiently guide updates in neural activity that eventually select the proper combination of modules for the task at hand.

In this work we have illustrated the learning mechanisms in systems with knowledge of only a handful of tasks. In real world settings, we likely enter any situation with a vast repertoire of modularized skills available to apply. On the one hand, this means that we very rarely have to learn new sub-skills from scratch; on the other, the number of possible recombinations might be very large. That said, there is also a vast amount of sensory, social, and linguistic information which can inform exactly what subset of modules is likely to be relevant [37, 48]. This leads to a very general view of how learning proceeds: upon entering a novel situation, executive systems integrate information about current goals and context to narrow the set of potentially useful modules, and compositional learning occurs in this narrowed space, leading to adaptive behavior. In a world built by and for humans inferring context is often automatic (from e.g. built environment, linguistic and social cues etc.). By contrast, the need to shape animal behavior in experimental settings can be interpreted as teaching the animal which modules are relevant for the current task [49–52]. Animals still come into the lab with a vast range of modular skills, but little context for how to apply them in the unnatural world of the laboratory. Shaping narrows the relevant set of skills to something low-dimensional (corresponding to low-rank task embeddings in our model) which allows fast and flexible performance in the experiment. Biologically, this process could be achieved by selectively inhibiting irrelevant dimensions, a mechanism likely localized in frontal regions such as the lateral prefrontal cortex (lPFC). Concurrently, reward-encoding areas could drive targeted weight adjustments aligned with the underlying compositional structure. These synaptic changes would subsequently modulate frontal activity to recruit the appropriate modules. Whatever the case, the learning advantages afforded by structural composition make it appealing from an evolutionary standpoint, and the neural evidence points towards the fact that biological brains implement this strategy.

We’d again like to emphasize that the structure in weights emerges naturally from training on related tasks, and this requires very few architectural specifications (mainly a separate population of embedding weights and the right initialization, or possibly the right regularization scheme). Once the system has reduced control to a low-dimensional space, then learning can be fast. The specific algorithms proposed in this work are examples of how the system can constrain learning to these spaces with minimal additional computations, all of which are self-supervised. There is likely a family of implementations that take take advantage of this structure, but they will all learning quickly by constraining learning using this structure. Therefore, the central prediction of this model is that during learning in compositional settings, the evolution of activity in areas responsible for encoding which modules to recruit should be low-dimensional.

Our analysis of the experiment in Tafazoli et al. show that this kind of structural learning is compatible with behavior and neural data, while models based on inference or less constrained learning are not. Though there are parts behavior we do not model explicitly (mainly the effects introduced by information in the sequencing of blocks) we nonetheless capture the important features of within block learning, where animals must discover which modules to combine. In particular, capturing both rapid acquisition of response axis and slower learning of the relevant feature required iterative learning within a neural subspace aligned to these relevant task features. Moreover, because our models are fundamentally neural models of computation, we were able to validate the compositional mechanisms in our model by comparing against neural data. Again, we found that structural composition was the only model which could reproduce the patterns of module re-engagement that were observed in data. In this way, modeling tasks with structural composition can serve as a kind of bridge between cognitive and neural domains. In a cognitive model, doing inference over an experimenter defined variable like the relevant feature (color vs. shape) can be directly translated as moving activity along the corresponding dimension in embedding space. Crucially, these dimensions in neural space emerge naturally as a result of learning multiple compositionally related tasks.

From a broader perspective, one can conceive of weak and strong structural learning as residing in a larger hierarchy of learning types that trade off restrictions on what kinds of target tasks can be learned with the amount of experience required to learn them. At one end, networks and animals can achieve strong 0-shot performance given instructions for the task, including in the form of natural language [22, 53]. Here no experience is required, but the only task acquired in this manner is the one being instructed. Strong structural composition can learn tasks that are within the inclusion/exclusion structure of the pre-training task set (as noted before, this setting applies to a vast range of experimental and real world settings [29, 30, 53–58]), and only require a handful of trials to achieve strong performance. Weak structural composition can learn tasks where modular components are combined or interpolated in the same low-dimensional subspace, and can learn efficiently with many dozens of trials. Modular composition can learn a target task that is made up of modules acquired during pre-training combined in any arbitrary manner, but learning may take many hundreds of trials [21, 59]. Finally, more traditional learning schemes that make updates to all of the weights of the system can learn any input-output mapping. But this can take thousands of trials, especially when learning from reward feedback, and can cause previous tasks to be forgotten. There is a rich literature of behavioral modeling that examines fast structural learning through inference vs. incremental learning via synaptic weight updates [46, 60–63]. One of the key insights of this work is that structure is not a binary. There are gradations of structure that systems can take advantage of in order to learn. The tradeoffs of instructed, strong, weak and modular composition followed by traditional weight updates neatly illustrate a widely held intuition: to learn quickly the system must making increasingly strong assumptions about how tasks are organized [49]. How biological agents gain the skills and experience required to form modules remains an open question. Theoretical work suggests that biological constraints may lead to regularization that has similar effect as small weight initializations [64]. Animal may also acquire this knowledge over many different timescales: evolutionary [49, 65], learning over a life time [66], or shaping in the laboratory [50–52].

Past theoretical work has studied systems similar to the linear compositional networks we used to derive our analytical results [26, 67]. In particular [23] investigated a nonlinear version of the same basic student-teacher set up and showed that learned module weights in the student must be a linear combination of module weights in the teacher. While that work mainly focused on the emergence of modules, our analysis demonstrates how and when the structure of module usage is encoded in weights at the end of learning. Other recent theoretical work has examined how parallelogram (or “abstract”) geometry emerges in networks trained on multi-dimensional binary classification tasks [68]. Networks had no explicit notion of modules, nor a code for module usage across tasks. Rather, they received random inputs and had to match each of the inputs to a multidimensional binary vector in the output which indicated which of multiple classes the input belonged to. Under these conditions parallelogram representations emerged, matching the binary code in the output. Our theory and subsequent simulations show that for a parallelogram geometry to emerge, it is sufficient for the structure of module usage across the task set (encoded in **G**^∗^) to be binary.

Since some of the first work on neural compositionality, these abstract, parallelogram geometries have played a central role. Such organization emerges in both multitasking recurrent and feedforward network models [22, 25, 69, 70]. In neural recordings, one of the most common proxy measures of this geometry is the cross-condition decoding used in Fig. 7. High cross decoding accuracy has been observed in animals and humans performing tasks with overlapping demands [33, 34, 40, 71]. Intriguingly, [34] observed that if neural representations did form a consistent geometry then humans quickly switched responses after a change in task demands. In this work we offer a precise mathematical description of what geometry should emerge given the organization of the task set. This allows us to predict cross-conditional decoding for any compositional task set organization prior to training a model. Given the rise of multitasking paradigms and the prevalence of cross-conditional decoding in neural data analysis, such predictions could prove valuable in designing experiments and explaining their results.

While the importance of abstract geometries for flexible cognition and learning has been recognized since these patterns were first identified [33], our work provides the first mechanistic model describing how rapid adaptation depends on this structure. This was possible because our analyzes showed exactly how structure in the *weights* matches structure in the task set. This in turn revealed the small number of directions in weight space that effectively control the entire system. Without the formal expressions in Eqs. 8a–9b, the presence of representational geometry in population activity remains highly suggestive, but offers no clear mechanism for updating activity in a way that adheres to this underlying organization. Finally, our work also shows that this abstract parallelogram geometry is an important special case of a much more general framework for predicting the geometry of representations in non-binary settings, as demonstrated in Fig. 3f-g. Indeed, this framework may explain work which showed motor learning is significantly faster for tasks parameterized by continuous variables on a low-dimensional manifold than for tasks with independent parameters [72].

The requirement that compositional systems possess a set of rules that govern module recombination has elsewhere been referred to as the syntactic dimension of composition, in rough analogy with the rules that specify valid linguistic utterances. Structural composition as defined here is but one relatively simple scheme for constraining module recombination in neural systems. Future work can build on the understanding we’ve developed in this setting to probe the neural implementation of syntactic composition in increasingly complicated settings, including sequences, predictive tasks, navigation, and eventually natural language [28, 73–75].

Though there are many open questions in this direction, overall we believe this work emphasizes the fundamental importance of the syntactic aspect of composition. Having modularized skills is certainly advantageous. But when the system has both modularized skills and a representation of how those modules can be recombined, then learning can become very fast. Specifying how exactly these rules are represented by neural systems also leads to a more unified picture of experimental and simulation results: observed geometries fall directly out of the structure of the task set, and learning is effected by this geometry because the geometry itself constrains how the system will learn. Our hope is that examining how neurons represent syntax will lead to an even more coherent view of compositional mechanisms in increasingly sophisticated task domains, and eventually help us understand the incredible fluidity of our own cognitive abilities.

## Supporting information

Mathematical Supplement

## Methods

### Task Classes

In the text we test five main classes of tasks, Student-Teacher Tasks, Wave Tasks, Go Tasks, ContinuousGo Tasks, and Decision Making Tasks. These basic task classes can be used to construct different task sets. Crucially, though different classes require fundamentally different computations, we can construct task sets such that they are made using same compositional structure. First we describe the Student-Teacher class and the formula for task set construction in this context. This will serve as the template for task set construction in more complex classes.

#### Student-Teacher Task Class

For this case we consider a bi-linear model, with the teacher given in Eq. (2) and the student in Eq. (6). The input, **x**, and output, **y**_*τ*_ are both 64-dimensional, so the weights are 64 × 64

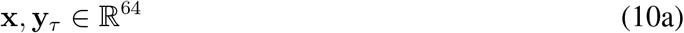

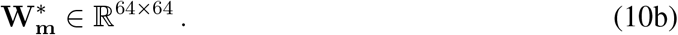

Both the components of **x** and the components of the weights, 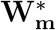 unit variance Gaussian, are drawn from a zero mean,

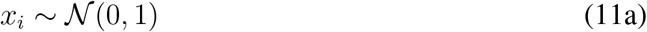

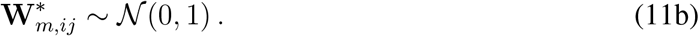

The elements of the matrix **G**^∗^ determine the extent to which module *m* contributes to the output on task *τ* ; the value of *τ* is just an integer which specifies the current task and hence the relevant row of **G**^∗^. We studied five task sets defined by the following matrices **G**^∗^,

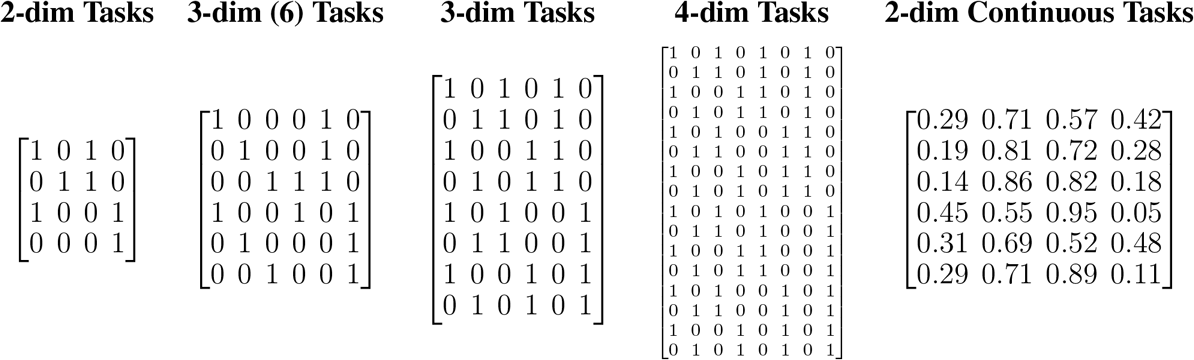

The main text primarily deals with 3-dim tasks and 2-dim Continuous Tasks; the remainder of the task sets are treated in the Supplementary Information.

#### Wave Task Class

For our wave task set there is no input and data points consist of

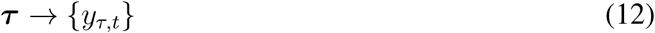

where *τ* is a one-hot vector (with a 1 at index *τ*) and the set {*y*_*τ,t*_} is a target time-series made of multiple single period cosine waves concatenated along the time dimension. Individual cosine waves each have a different period and play the role of modules in the previous task. As above, each task is defined by what modules are included in producing target outputs. For a given period *p* the targets for that period are given by

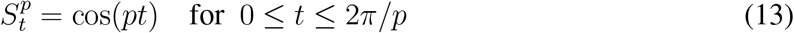

The periods we used were

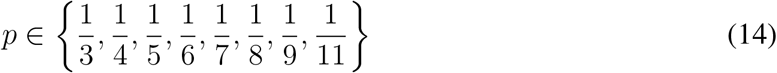

So, for example, the 2-D wave task set uses 4 periods in total 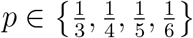. Target outputs for the first task *τ* = 1 are then are produced using periods 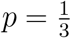 and 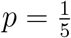. The overall target is given by

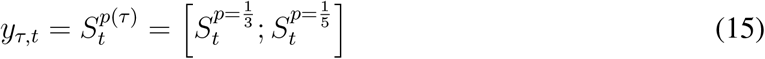

where brackets denote concatenation along the time dimension. We don’t construct a continuous task set for the wave class. However, one can build an analog to all other task sets by taking the first *M* periods and using the rows of the desired **G**^∗^ matrix as a binary mask which determines which periods are included in constructing the overall target output. Although we could’ve used any four periods as the components of our task set, to be concrete, for the 2-D task set all target outputs are constructed as follows,

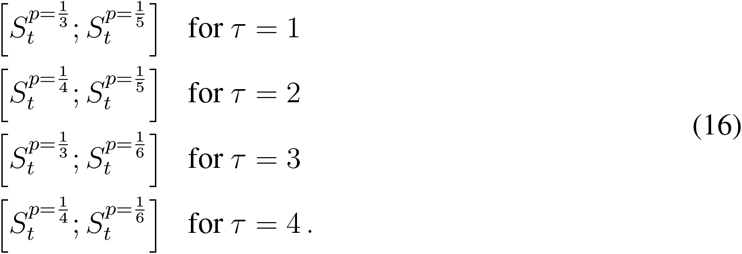

In all simulations we use a step size of *dt* = 1.0.

For the waves class reward is given by the mean number of time steps where the squared difference between the response and the target is less than a threshold *δ* = 0.05

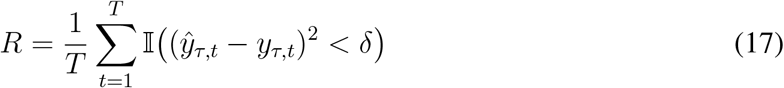

where I(·) is the indicated function which takes a value of 1 if its argument is true and 0 otherwise.

#### Go Task Classes

Both the Go task class and the Decision-Making (DM) task class are taken from previous work on multitasking RNNs [21, 25]. At a high level, the Go class involves sensory inputs from multiple input modalities and motor responses in one out of multiple possible output modalities. Tasks are built by designating one relevant input modality, one relevant output modality, and whether networks must respond in the same of the opposite direction of the relevant input (‘Go’ vs. ‘AntiGo’). As above, each trial includes a task cue *τ* . Trials unfold over time and consists of three trial periods. Task information, *τ* is always present in the input. In the preparatory period, the model receives only task information, *τ*, and no sensory input. During this period the model must also maintain fixation by outputting high activity in the fixation output unit. In the stimulus period, the sensory input is present and the model must also maintain fixation. In the response period sensory input is absent and the model can break fixation and report its response. A given trial consists of

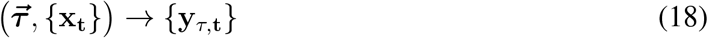

where {**x**_*t*_} is the time series of sensory inputs and {**y**_*τ,t*_} is a time series of target outputs for a given task.

The sensory input, **x**^stim^, consists of information about each modality along with a cue that tells participants whether to fixate or respond. The information about modality *i* consists of a two dimensional vector that lies on a circle,

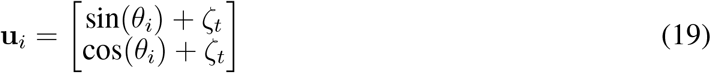

where *i* ranges from 1 to the total number of input modalities denoted ℐ, and *ζ* is a small amount of noise drawn in independently at every time step (*ζ*_*t*_ ∼ *N*(0, 0.1)). The cue is simply a binary variable, *u*_fix_ ∈ {0, 1}, where *u*_fix_ = 1 tells the subjects to fixate and *u*_fix_ = 0 tells the subjects to respond. The stimulus, **x**^stim^ is, then, a concatenation of the *u*_fix_ and **u**_*i*_,

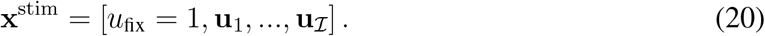

With this definition, **x**^stim^ ∈ ℝ^1+2ℐ^ . The other task periods consist of fixation unit followed by vectors of zeros where the stimulus information would appear.

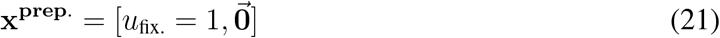

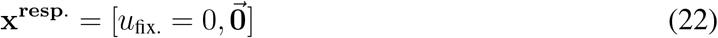

For all tasks, the preparatory period lasts for 10 time steps, the stimulus period lasts for 30 and the response period lasts for 10.

Target responses are built in a similar way. Like inputs, each response modality **v**_*i*_ is a two dimensional vector that encodes a 1-dimensional circular variable. Suppose for a given task *τ* the relevant input modality is *i*. If the task *τ* requires a standard Go response then the target angle is 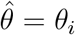; if the task requires an Anti-Go response then the target angle is 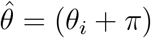. Then, if the relevant output modality is *j*, the target output is

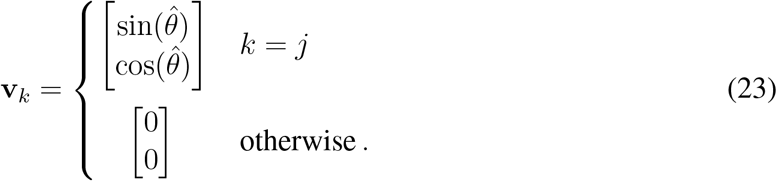

The overall target for the response period includes a target fixation unit, *v*_fix_. This should be set to 0 during the response period to represent a break in fixation and 0.95 during stimulus and preparatory periods to indicate the participant is fixating. Letting *ρ* be the total number of output modalities, we have

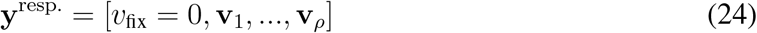

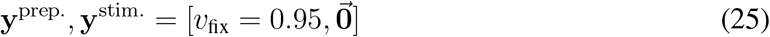

The structure of task sets in the Go class are determined by the relevant input modalities, the relevant response modalities, and the choice between a Go and an AntiGo target. With these components we can construct task sets that are analogous to 2-D Tasks, 3-D (6) tasks, and 3-D tasks introduced in the preceding sections. For 2-dim Tasks the total set of task components is {Mod1, Mod2, Go, AntiGo}, where Mod*i* denotes the relevant input modality (in this case we use only a single output modality). For the 3-dim (6) tasks this set in {Mod1, Mod2, Mod3, Go, AntiGo}. For 3-dim tasks {Mod1, Mod2, Go, AntiGo, Resp1, Resp2}, where Resp*i* denotes the relevant output modality. Like in the wave task, the demands for each task in a given task set can be obtained by taking these tasks and applying the rows of **G**^∗^ as a binary mask to the components of these sets.

Reward for this task class is binary and requires response to maintain fixation during the preparatory and stimulus periods and then respond within a threshold of the target angle. For convenience we’ll denote all the output channels excluding the fixation channel as **y**_**out**_ and **y**_*i*_ as the output channels corresponding to the *i*^*th*^ output modality. Here we set the threshold 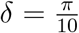 and recall that the target output modality is indexed by *j*

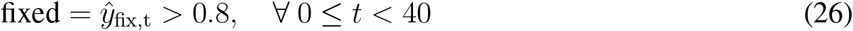

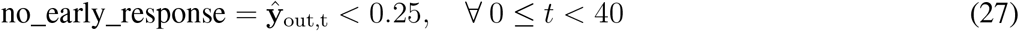

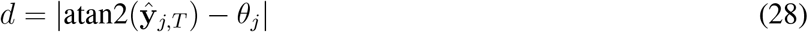

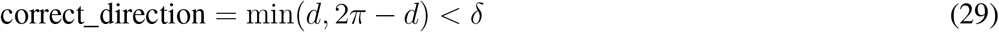

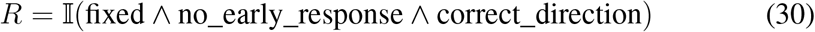

#### Continuous Go Task Class

We can construct an analog of the 2-dim continuous task set by defining task targets as a smooth interpolations between angles presented in two inputs modalities. For this task class there are exactly 2 inputs modalities and a single output modality. The task is constructed using two parameters, *α*_sen._ and *α*_resp_. Input information is generated in the same way as above. Given stimulus angles *θ*_1_ and *θ*_2_, target angles are generated taking *θ*^*′*^ = (1 − *α*_sen._)*θ*_1_ + *α*_sen._*θ*_2_, and 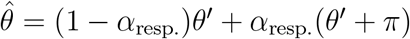. Target activations for units are then given by the above formula for *v*_1_. One can then construct the analogous task set to the 2-dim continuous task defined above by setting the values of (1 − *α*_sen._), *α*_sen_, (1 − *α*_resp._), *α*_resp._ to the values in the columns of the corresponding *G*^∗^. Rewards criteria for this task set is the same as the Go task class (Eq. 30).

#### DM Task Class

The Decision-Making task class shares many of the same features as the Go task class. It consists of a number of input modalities each encoding a 1-dimensional circular variable, and a number of target output modalities where the responses are reported.

In this case however there are two stimulus periods interspersed with a delay period. In each stimulus period, a different direction is presented. Each directional stimulus also has a different amplitude. Targets are generated by selecting the relevant input modality, selecting the direction of either the greatest (Strong) or the weakest (Weak) amplitude, and then reporting this direction in the correct response modality. Concretely, let 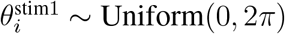 be the stimuli presented in the *i*^*th*^ input modality in the first stimulus period. The second stimulus direction 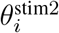 is drawn some offset from 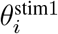

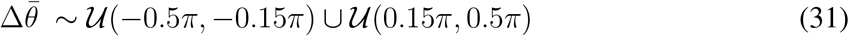

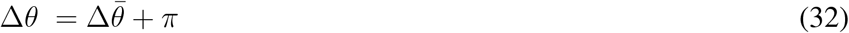

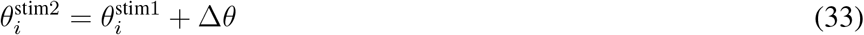

We draw a baseline amplitude *A*_base_ ∼ Uniform(0.5, 1.5) and a coherence coh from the set {− 0.2, −0.15, −0.1, 0.1, 0.15, 2.0}. Then inputs for stimulus period *k* in input modality *i* is given by

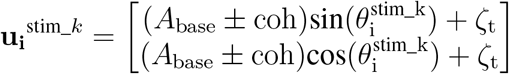

where *±* takes the sum if *k* = 1 and the minus if *k* = 2. Like above *ζ* is a small amount of noise drawn independently at every time step *ζ*_*t*_ ∼ *N*(0, 0.1) The target direction for a task requiring response for the strong direction (Strong) task with the *i* the relevant input modality is

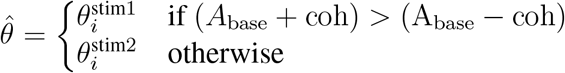

In the case of a Weak task the *>* sign in the condition is replaced by <. Overall input feature vectors **x**^**prep**.^, **x**^**stim**.^ and **x**^**resp**.^ and output features **y**^**prep**.^, **y**^**stim**.^ and **y**^**resp**.^ are constructed in the same way as the Go task. The input during the delay **x**^**delay**^ = **x**^**prep**.^. For these tasks the period lengths are 10, 10, 6, 10, 4, 10 for the preparatory, stimulus 1, delay, stimulus 2, delay and response periods respectively. Again, the structure of task sets is determined by the relevant inputs modalities, the relevant response modalities, and the choice between Strong and Weak decision making types. For 2-dim tasks the set of task components is {Mod1, Mod2, Strong, Weak} (in this case we use only a single output modality). For the 3-dim (6) tasks the set is {Mod1, Mod2, Mod3, Strong, Weak} (again with a single output modality). For 3-dim tasks the set is {Mod1, Mod2, Strong, Weak, Resp1, Resp2}. The demands for each task in a given task set is obtained by applying the rows of **G**^∗^ as a binary mask to the components of these sets. Reward is delivered in the same way as the Go task class.

#### Tafazoli et al. Task Class

We also build a model version of the task set used in [29]. In this experiment, participants receive a single vision stimulus per trial, which has both a color feature (some shade between red and green) and a shape feature (shape is morphed between a tee shape and a bunny shape, see Fig. 6a. in the main text). Participants then had to decide what category the stimuli belong to based on what the relevant stimulus feature was on that trial. For example, in a color trial participants decided whether the stimuli is more green or more red. Participants then had to report their response using the proper response axis. Response axis 1 consists of pressing on the upper left and bottom right regions of the screen while response axis 2 requires press on upper right and bottom left. Each individual task in the task set combines these two components e.g. the shape, response axis 1 task (S1) requires the participant to categorize the stimuli based on its shape feature and response using the response axis 1.

To model this task, we simply represent the features of a stimuli as two circular variables (one for color and one for shape) which serve as sensory input. In particular for each trial, we draw *θ*_color_, 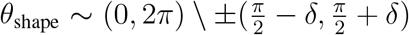 i.e. drawn uniformly from the circle except for a small region around the decision boundary at 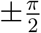. For both features, if 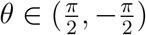 then it belongs to category 1 and if 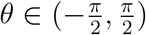 then the stimulus belongs to category 2.

Like in previous tasks there are three task periods, a preparatory period, stimulus period, and response period. Again, we will represent each circular variable in terms of their sin and cos components. However, in this case there is no fixation cue. The input vector for this task is 4 dimensional. There is no stimulus input during the preparatory period

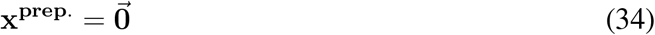

For each time point in the stimulus period, noisy stimulus values are drawn from von Mises distributions centered on *θ*_color_ and *θ*_shape_ with concentration parameters *κ*_color_ and *κ*_shape_ respectively.

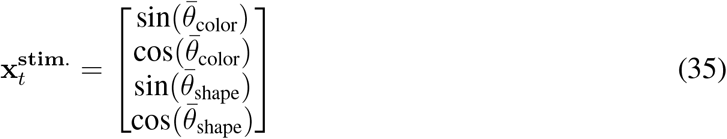

where 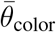 and 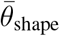 are given by

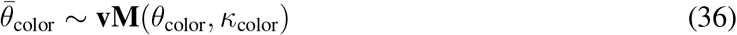

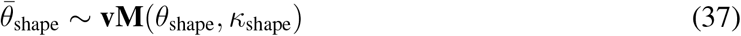

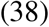

Finally, there is also no stimulus input during the response period.

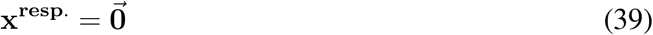

Across all tasks the preparatory period is 10 time steps, the stimulus period is 20 time steps and the response period is 10 time steps.

During the response period, target outputs are given by a two dimensional vector where each dimension corresponds to one of the response axes. If the task requires a response along the first axis then that component is ±1 depending on the category of the relevant stimulus feature and the alternative vector component remains 0. Formally, let

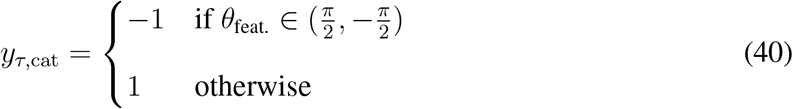

where *θ*_feat._ is the angle of the relevant stimulus feature for task *τ* . Then the response vector is given by

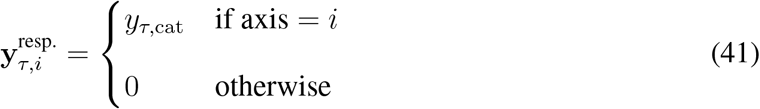

During the preparatory and stimulus periods the target output is the zero vector.

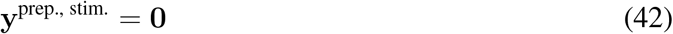

### Neural Network Architectures and Initializations

#### Student Teacher Models

All student models tested in simulation closely emulate the architecture of the teacher. In all cases, student modules

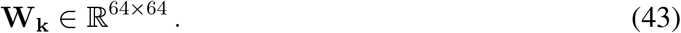

Unless otherwise specified in the main text we also set *M* = *K*; i.e., the number of modules in the student is the same as the total in the teacher. For small weight initializations each student module weight

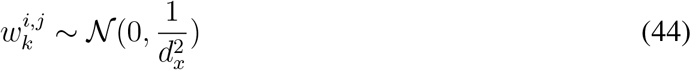

where *d*_*x*_ is the dimension of the input data; in all simulations *d*_*x*_ = 10. For

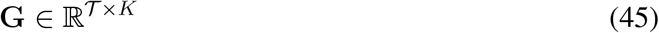

small weight initialization is given by

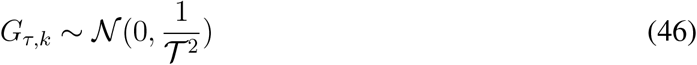

where *T* is the number of tasks.

For large weight initialization

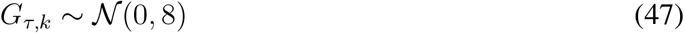

and initialization for **W**_*k*_ was kept the same.

#### Recurrent Networks

All RNNs that trained on the Wave, Decision-Making, Go, and ContinuousGo task classes are implemented as Gated Recurrent Units (GRUs) [77]. In each case models produce a task embedding for task *τ* by passing a one-hot vector 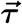 through a set of embedding weights 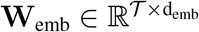. The resulting embedding vector **emb** is then concatenated with the sensory stimulus **x**_**t**_ at every time step. This concatenation is fed to the RNN as input. The state of the RNN, **h**_*t*_ is passed through a set of readout weights *W*_out_ and a nonlinearity *ϕ* to produce an output **ŷ**_*t*_.

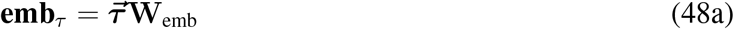

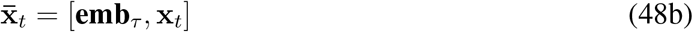

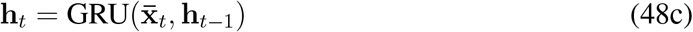

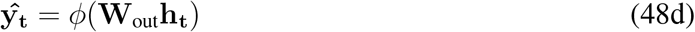

where here we’ve used 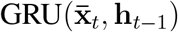 as a shorthand for the following GRU update equations given input vector 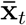 and previous state **h**_*t*−1_.

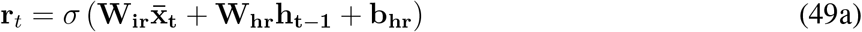

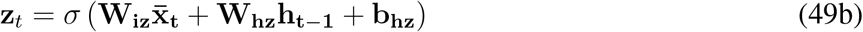

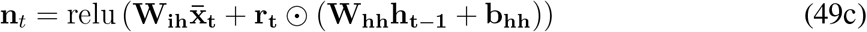

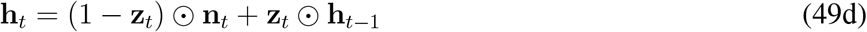

In the main text, where we say that we “freeze RNN weights” we mean the set of parameters **W**_**ir**_, **W**_**hr**_, **W**_**iz**_, **W**_**hz**_, **W**_**ih**_, **W**_**hh**_, **b**_**hr**_, **b**_**hz**_, **b**_**hh**_ . For all models we set the initial state of the RNN to **h**_0_ = **0**. All RNNs use a ReLU non-linearity. Let *d*_*h*_ be the dimension of the RNN state. Then both small weight initialization are given by the Table 1, where **W**_**h**_,**b**_**h**_ and **W**_**in**_ refer to weights sub-scripted with h and i respectively in Eq 49.

**Table 1:** Comparison of Small and Large Weight Initializations for the RNN.

| Weight | Small Weight Initialization | Large Weight Initialization |
| --- | --- | --- |
| Embedding Weights ($\mathbf{W}_{\text{emb}}$) | $w_{\text{emb}}^{i,j} \sim \mathcal{N}\left(0, \frac{1}{d_{\text{emb}}^2}\right)$ | $w_{\text{emb}}^{i,j} \sim \mathcal{N}(0, 1)$ |
| Recurrent Weights ($\mathbf{W}_{\text{h}}$) | $w_h^{i,j} \sim \mathcal{N}\left(0, \frac{1}{d_h^2}\right)$ | $\mathbf{W}_{\text{h}} = I$ |
| Input Weights ($\mathbf{W}_{\text{in}}$) | $w_{\text{in}}^{i,j} \sim \mathcal{N}\left(0, \frac{1}{d_h^2}\right)$ | $w_{\text{in}}^{i,j} \sim \mathcal{N}\left(0, \frac{1}{d_h}\right)$ |
| Bias ($\mathbf{b}_{\text{h}}$) | $b_i \sim \mathcal{N}\left(0, \frac{1}{d_h^2}\right)$ | $b_i = 0$ |
| Output Weights ($\mathbf{W}_{\text{out}}$) | $w_{\text{out}}^{i,j} \sim \mathbb{U}\left(-\sqrt{\frac{1}{d_h}}, \sqrt{\frac{1}{d_h}}\right)$ | $w_{\text{out}}^{i,j} \sim \mathbb{U}\left(-\sqrt{\frac{1}{d_h}}, \sqrt{\frac{1}{d_h}}\right)$ |

**Table 2:** Free parameters for each model.

| Model | Free Parameters |
| --- | --- |
| Inference | $\kappa_{\text{shape}}, \kappa_{\text{color}}, \lambda_{\text{reg}}, \alpha_b, w_0, w_1, w_2, \tilde{w}_0, \tilde{w}_1, \tilde{w}_2$ (10) |
| Modular | $\kappa_{\text{shape}}, \kappa_{\text{color}}, \lambda_{\text{reg}}, \eta_{\mu}, \eta_{\sigma}, \sigma_0, \tilde{\sigma}_0, \mu_0^{\text{feat.}}, \mu_0^{\text{axis.}}, \tilde{\mu}_0^{\text{feat.}}, \tilde{\mu}_0^{\text{axis.}}$ (11) |
| Structural | $\kappa_{\text{shape}}, \kappa_{\text{color}}, \lambda_{\text{reg}}, \eta_{\text{feat.}}, \eta_{\text{resp.}}, \mu_0^{\text{feat.}}, \mu_0^{\text{axis.}}, \nu_0^{\text{feat.}}, \nu_0^{\text{axis.}}, \tilde{\mu}_0^{\text{feat.}}, \tilde{\mu}_0^{\text{axis.}}, \tilde{\nu}_0^{\text{feat.}}, \tilde{\nu}_0^{\text{axis.}}$ (13) |

For the Wave task class, and Tafazoli et al. task class, *d*_*h*_ = 128, *d*_emb_ = 64. For the wave task class *ϕ* was simply the identity mapping i.e. the readout is a linear mapping of RNN states. Tafazoli et al. task class uses *ϕ* = tanh(). For Decision-Making, Go, and ContinuousGo task classes, *d*_*h*_ = 512, *d*_emb_ = 128 and *ϕ* = tanh().

### Neural Network Training

For all models we minimized mean squared error using SGD with the Adam optimizer. All models are implemented and trained using PyTorch.

#### Student Training

Student networks were trained using stochastic gradient descent on the mean squared error (MSE) loss between student output *ŷ* and the target output *y* produce by the teacher network. During training we used batch size of 32 and a learning rate of *η* = 1*e*^−2^. In total, we trained each network for 50,000 batches.

#### RNN Training

For the Wave, Decision-Making, Go and ContinuousGo task classes, RRNs are trained to minimize the MSE loss between model output *ŷ*_*t*_ and the target output *y*_*t*_, averaged over the time. For a given batch

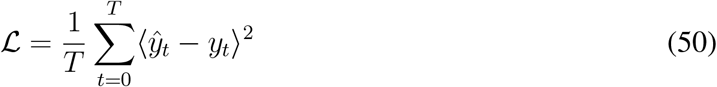

In all cases, we perform gradient descent with an Adam optimizer [78]. For both the Wave task class and Tafazoli et al. task class, we train for a total of 50,000 batches using an initial learning rate of *η* = 1*e*^−3^, and batch size of 64. In the Tafazoli et al. task class we set *κ*_color_, *κ*_shape_ = 5.0 during training. For the Decision-Making, Go, and ContinuousGo task classes we train for 100,000 batches using an initial learning rate of *η* = 1*e*^−3^ and a batch size of 32. For all RNN tasks we also decay the learning rate by a factor of 0.97 every 1000 batches. Results presented in the main text are robust to a range of training hyperparameter choices, and are primarily effected by the weight initialization.

### RNN Modularity Analysis

To illustrate the modularity of computations in multitasking RNNs (Fig. 1c), we plot the time averaged, normalized variance of each unit across tasks [25]. In Fig. 1c the network was trained on the 3-D Decision-Making task set with Mod1-Strong-Resp1 held out of training. Then, for each task, we compute the variance across time for each unit using 100 trials, and for each unit we normalize by the maximum variance over tasks. For the held out task we cue the network with embedding activity extracted from the missing vertex of the underlying parallelogram structure (the blue x in Fig. 1e; see Compositional Learning from Reinforcement below). Subsequently, we perform k-means clustering by treating the normed variance of each unit as a feature vector over tasks. We test values of k between 2 and 30, and we choose the cluster whose k has the maximum silhouette score.

Note that while previous studies performed this analysis for individual trial periods (e.g. clustered activity only during the stimulus period), this was primarily important for identifying clusters of units in networks trained on tasks with very different timing structures (e.g. networks trained on tasks with both a single stimulus, and with two stimuli interspersed with a delay) [21]. In our networks, the structure of timing within a task set is consistent, so analyzing per trial period is not necessary to identify the important modular structure in our networks.

### Empirical Tests of Compositional Structure

To test the conjecture given in Eqs. (9a) and (9b) in the main text, we need to compute the rank and optimal rotation of 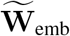. To measure the effective rank of 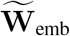 after training we simply plot the cumulative proportion of variance explained by each successive singular value *s*_*i*_,

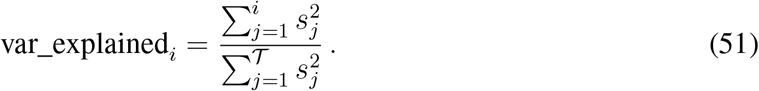

The effective rank of 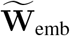 is defined as the minimum *i* such that var_explained_*i*_ *>* 0.9. To test if **Ũ**_emb_ (left singular vectors of the mean centered weight matrix 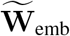) is a rotation (and possibly a reflection) away from 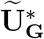, we first find the optimal rotation *R* between the two matrices and measure the root mean squared between the resulting set of points. Finding involves solving the orthogonal Procrustes problem [81] To do that we use scipy.linalg.orthogonal_procrustes. Importantly, in each case *ℛ* ∈ *ℝ* ^*r×r*^ i.e. a rotation in the rank of the matrix **G**^∗^ that encodes the task set structure, rather than the number of tasks *T* . We then take

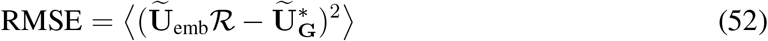

Where 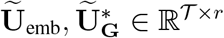.

For every task set we build a set of control weights weights 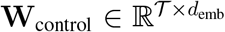, where 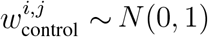. We draw 25 such matrices and extract the RMSE and variance explained as outlined above. Note that for both the control weights and weights of models initialized with large weights the effective rank is generally greater than the rank of 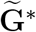. Despite this for all of our rotation measures and visualizations we only take the first *r* columns of the left singular vectors **Ũ**_emb_ for these controls. Hence, the error measures reported in the main text and Supplementary Information for these cases are conservative; they describe error only in the primary components of these matrices while neglecting a substantial portion of the remaining variance.

To assess statistical significance of RMSE plotted in Fig. 3d., f. we performed a Mann-Whitney U-test between the 5 randomly initialized models for each task class and the distribution of 25 control matrices, and then perform Bonferroni correction to account for multiple comparisons.

### Compositional Learning from Reinforcement

Suppose we begin with an RNN that has been pre-trained on a given task set with a particular task, *τ*_*h*.*o*._, held out of training. Our goal is to learn the task embedding vector for the held out task using only reward feedback provided on a trial-by-trial basis. We allow the network to learn over a run of trials *i* 1…*N* . To simplify, let 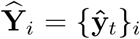 be the set of model outputs over time in trial *i*, **X**_*i*_ = **x**_*t i*_ the set of stimulus inputs over time in trial *i*, and *ρ*_*h*.*o*_ the reward function for the held out task (reward criteria are given in Task Classes section). As a short hand we’ll write the output of the RNN (Eq. 48) as a function of the stimulus inputs and embedding vector on trial *i*,

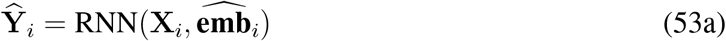

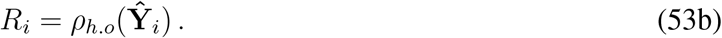

We assume the weights of the RNN are frozen, and adjust 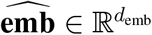 to maximize reward.

#### Modular Learning

In the modular setting, we treat 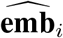 as a control signal parameterized as a multivariate normal distribution,

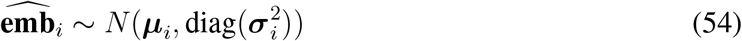

where ***µ*** is a vector of means and the covariance is a diagonal matrix built from the vector ***σ***^2^. We update activity in this control signal according to the REINFORCE algorithm. Updates are given by

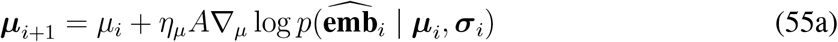

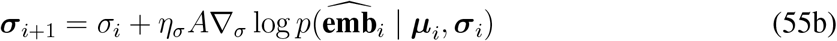

where *A* = *R*_*i*_ − *b*_*i*_ is the advantage score. Here *b*_*t*_ is a reward baseline term with momentum *υ* that is updated according to

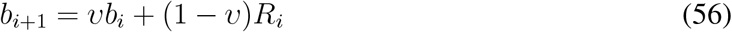

In all cases we set *υ* = 0.9 and *b*_0_ = 0. We set ***µ***_0_ = ⟨**W**_emb_⟩ where the angle brackets indicate an average over all task embeddings seen during pre-training, and ***σ***_0_ = *σ*_init_, where *σ*_init_ is a hyperparameter. We also set a maximum possible variance for learning, so after each update we enforce ***σ***_*i*_ = min(***σ***_*i*_, *σ*_max_) where *σ*_max_ is also treated as a hyperparameter. Finally, both learning rates *η*_*µ*_ and *η*_*σ*_ are hyperparameters of learning. Details on hyperparameter search is given below.

#### Structural Composition Mapping

To take advantage of the compositional structure, we can use the fact that the embedding weights, **W**_emb_, live in an *r*-dimensional space. In both strong and weak structural composition this will allow us to learn the embedding activity for the held out task using a low-dimensional control signal **c** ∈ *ℝ*^*r*^. We first define a function *F* that maps from 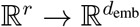 in the basis defined by 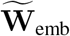

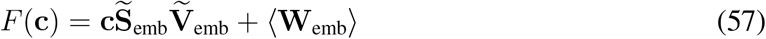

where 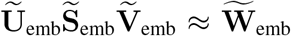 is the truncated singular value decomposition (we only keep the number of singular values and components required to capture 90% of the variance in 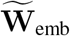).

#### Weak Structural Learning

In this setting our goal is to update a continuous vector of activity **c** ∈ ℝ^*r*^ to learn the held out task. Except for the fact that we are now learning over a lower dimensional control signal, learning proceeds in exactly the same way as Modular Learning outlined above. We assume **c** is drawn from a multivariate normal

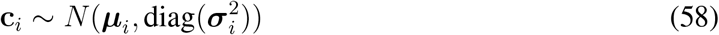

and update the parameters of the distribution according to the REINFORCE update

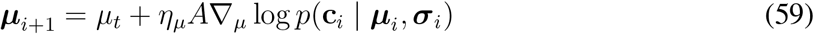

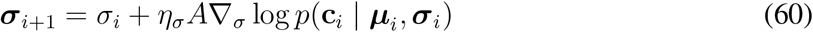

where *A* is the advantage score outlined in the Modular Learning section. The embedding vector activations that are fed to the pre-trained RNN are then given by

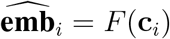

The set of hyperparameters for weak structural learning is the same as in modular learning. We set the initial mean by taking the average of pre-trained task embeddings and then projecting that into the low dimensional space i.e. ***µ***_0_ = *F* ^−1^(⟨**W**_emb_⟩).

#### Strong Structural Learning

For strong structural learning our goal is again to update a continuous, low-dimensional control signal. For binary tasks we show in the Mathematical Supplement Section 2 that the rows of *Ũ*_emb_ will form a subset of the vertices of an *r*-dimensional parallelogram. To take advantage of this geometric regularity, we introduce an affine transformation T from ℝ^*r*^→ ℝ^*r*^ that maps vertices on this *r*-dimensional parallelogram onto unit aligned *r*-dimensional hypercube. First we defined an anchor index ind_anchor_ which is defined as the task embedding with greatest cumulative similarity to other tasks (this simply ensures that we don’t choose an anchor index corresponding to a vertex adjacent to a held out vertex),

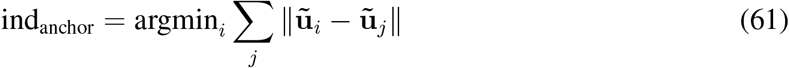

where **ũ**_*i*_ is the *i*^th^ row of **Ũ**_emb_. We’ll define 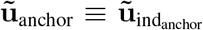. We then take the *r* closest **ũ**_*i*_ by euclidean distance and stack these row vectors into an (*r* + 1) *× r* matrix **Ũ**_corner_. Our scaling factor is given by *γ* = |**ũ**_anchor_ − **ũ**_*j*_|, where **ũ**_*j*_ is one of the *r* nearest neighbors to the anchor. Finally, we take the inverse 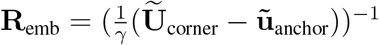, which essentially maps our scaled and shifted matrix onto the unit axes. We can then write our overall T as

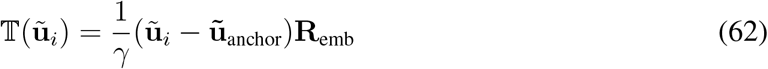

Given the regularities ensured by a binary **G**^∗^, T takes the parallelogram structure of **Ũ**_emb_ and maps it onto the *r* dimensional unit cube. In particular, for each **ũ**_*i*_ (i.e. the low-dimensional task representations), T(**ũ**_*i*_) ∈ {0, 1}^*r*^. Further, because of the geometric regularities in **Ũ**_emb_, the missing vertices which correspond to held out tasks will also lie on a vertex of the hypercube in the image of T. As we’ll see this will allow us to orient exploration around the vertices of the parallelogram and then map back to the original embedding space via T^−1^.

Since we are primarily interested in exploring the vertices of this hypercube during learning, we parameterize our control signal using a multidimensional Beta distribution, i.e.

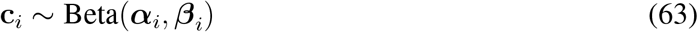

where the concentration vectors ***α, β*** are initialized with small values. This places the majority of the density around 0 and 1 along each dimension, which results in draws from the vertices of the hypercube. With small values for ***α, β*** the gradient of the likelihood becomes unstable and the REINFORCE update fails to learn in context of this strong prior. Instead, we formulate a heuristic which mimics REINFORCE’s tendency to move the distribution closer to control signal on rewarded trials and further away on unrewarded trials,

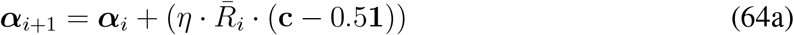

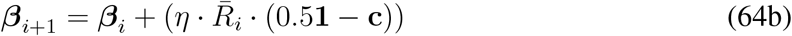

In this case we 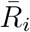 is the signed reward. Here we use

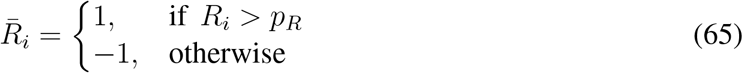

where *p*_*R*_ is the prior expectation of reward. For all task classes we set *p*_*R*_ = 0.95 except for wave tasks where *p*_*R*_ = 0.90.

At each trial we map to an embedding vector via

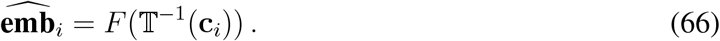

Hyperparameters for strong structural learning include the learning rate *η* and initial concentration value *a*_init_ such that 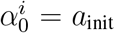 and 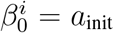. The method for selecting hyperparameters in given below.

For weak structural learning we can learn **c** in the space defined by basis of *W*_emb_, or in the space transformed by T. We found that learning in either space led to qualitative similar results. In the main text we report weak structural learning in the image of T because this was the fairest comparison to strong structural learning.

#### Hyperparameter Search

For each learning algorithm (modular learning, weak structural learning, and strong structural learning) we perform a grid search over the set of hyperparameters to determine which values lead to the fastest learning. The speed of learning is determined by the number of trials required for the model to reach performance criteria of 90% on the held out task. Performance is computed as the average over a sliding window of 10 trials. For each held out task, we test learning on a set of five models each with a different random initialization. For each of these models, we perform a set of *n*_*runs* independent learning runs each consisting of *N* trials where the model attempts to learn the task. The best set of hyperparameters for a given learning algorithm on a given model seed is the set which results in the minimum average trials to criteria over *n*_*runs*.

For modular learning, and weak structural learning *n*_*runs* = 5, *N* = 1000 and the grid of hyperparameters is given by

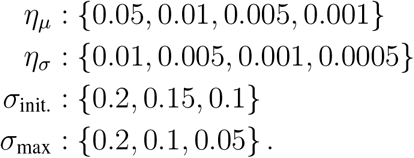

For strong structural learning *n*_*runs* = 10, *N* = 250 and the grid of hyperparameters is given by

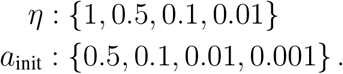

#### Statistical Tests for Learning

To asses statistical significance among different learning curves (indicated by bars above curves in Fig. 4d.-g.) across each type of learning we first performed an independent t-test between models at each time point. Consecutive significant points (p < 0.05) were grouped into clusters, with each cluster’s mass defined as the sum of its t-statistics. A null distribution of cluster masses was generated by randomly permuting learning type labels 5,000 times and taking the maximum mass for each permutation. Clusters were significant if their mass exceeded 95% of the null distribution. Only significant clusters of longer than 30 contiguous trials are plotted in Fig. 4.

To assess statistical significance of the number of trial required to reach 90% performance threshold across learning types, we performed a Mann-Whitney U-test between the distribution of learning runs and then Bonferroni correction to account for multiple comparisons.

### Model Fitting

We fit three different models to behavioral data from Tafazoli et al. [29]. Each attempts to match the time course of task discovery in blocks with true underlying tasks S1, C2, C1. First, we give a description of the models themselves and afterwards describe the fitting procedure used to tune parameters.

All models are based on the same set of RNNs (5 different random initializations using small weights), pretrained on S1, C2, C1 tasks. After training, when models have reached high performance on the task set, our modeling assumption is that the process of task discovery in a block is driven by changes in the task embedding vector, 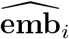, rather than recurrent weights. Hence, **W**_rec_, **W**_out_ and **W**_in_ are frozen throughout. Actions taken by the model at trial *i* are draws from a 4-dimensional categorical distribution (each dimension corresponding to one of the possible actions: bottom-left, top-left, top-right and bottom-right). At each trial *i*, the model produces a timeseries of outputs **Y**_*i*_ via Eq. 53a. For this task output is a 2-dimensional vector with outputs between 1 and -1. Flatten these outputs to a 2-dim vector with values normalized to 0 and 1. We then average the output over the last 5 times steps of each trial, resulting in a 4-dimensional vector **z**. Since the weights of the model are frozen, the action distribution on a trial by trial basis depends on sensory inputs **x**_*i*_ and embedding activity 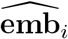. To emphasize this we subsume RNN equations 53a and output averaging into a single function and write

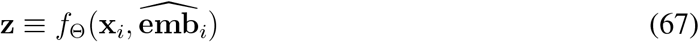

where Θ is the set of frozen model weights. In this case the action distribution is given by

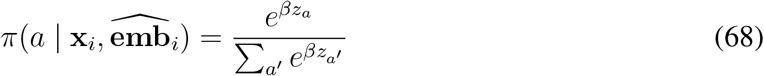

where *β* is the inverse temperature parameter. For all models we used a fixed *β* = 10.0 The action is then sampled from a categorical distribution:

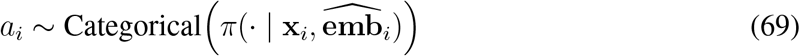

All models include free parameters *κ*_shape_ and *κ*_color_ which control the amount sensory noise applied to each aspect of the stimulus feature (see Task Classes).

The structure of the sequence of blocks (Fig. 6b., bottom) is organized so that following a C2 block there is always either an S1 or C1 block, and following an S1 or C1 block there is always a C2 block. This means that it is possible in principle for the animal to infer whether the upcoming block is a C2 block or either a S1 or C1 block, although animals take advantage of this information to varying degrees. We don’t try and model behavior that emerges from sequential information mechanistically, and instead focus on within block task discovery. Instead, to capture some of the sequential effects of block ordering, all models include a parameter *λ*_reg_ which regularizes the process of task discovery for blocks that follow C2 blocks (i.e. S1/C1 blocks). All models are also have prior parameters which determine the starting point of the policy for all C2 blocks and for all blocks after C2 blocks. Details are provided for each model below.

Models differ primarily in how they update embedding vector 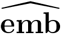, which in turn changes RNN dynamics and outputs **z**.

#### Inference Model

The inference model maintains a belief *b*_*i*_(*τ*) that *τ* is the true underlying task at trial *i* in the block. The model’s updated belief at the next time step is a linear interpolation between the current belief and the posterior belief inferred from the reward signal. The full posterior is given by

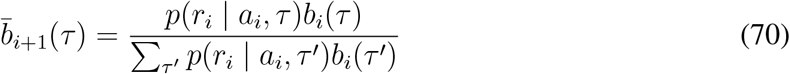

where the likelihood is determined by the probability of producing an action given a task embedding **emb**_*τ*_ and whether or not that action was rewarded

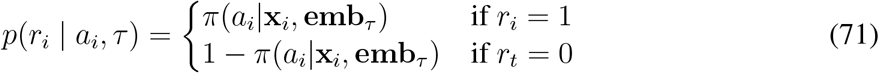

The the belief at the next trial is then a linear interpolation between the full posterior update and the belief at the previous time step.

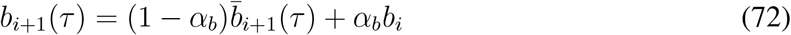

where *α*_*b*_ is a free parameter of the model.

In blocks S1/C1 (which always follow a C2 block), belief is regularized by interpolating with a uniform distribution U over the three tasks.

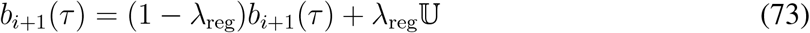

The task embedding vector at any trial *i* is obtained by drawing a task *τ* from **b**_*i*_ and setting the embedding on the current trial to the pretrained embedding for the corresponding task.

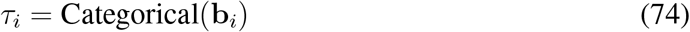

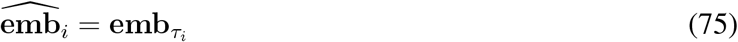

The prior belief over tasks at the start of each block is parameterized with weights (*w*_0_, *w*_1_, *w*_2_), normalized to sum to one:

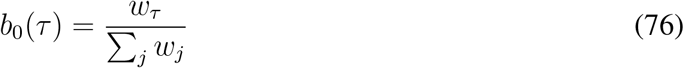

When the previous block is C2, a separate set of prior weights 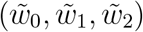 is used, allowing the model to capture any systematic bias in task beliefs following that specific rule context.

#### Modular Model

Update equations for the modular model follow those outlined in Weak Structural Learning. After each trial’s REINFORCE update, ***σ*** is clipped to a maximum of *σ*_max_ = 0.15. When the previous block was C2 (i.e. in S1/C1 blocks), ***σ*** is additionally mixed toward 1 (maximum uncertainty):

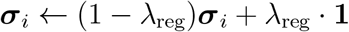

the learning rates *η*_*µ*_ and *η*_*σ*_ are free parameters.

At the start of each block, we use a initial standard deviation *σ*_0_ that is broadcast across all embedding dimensions:

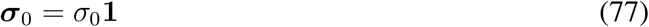

To get the initial mean vector *µ*_0_ we take advantage of the Structural Composition Mapping. We fit two scalars 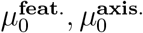, which gives the location of the initial embedding in terms of the relevant low-dimensional subspace, and then let

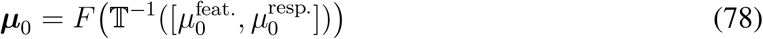

where *F* is defined in Eq. 57 and T is defined in Eq. 62. This allows us to efficiently fit the prior distribution of animals actions in a block using few parameters, and in a way that is comparable to the Inference Model and the Structural Model. This is crucial, as we mainly want to compare task discovery dynamics within a block. Note, however, that once task discovery commences there are no constrains which restrict the embedding to the low-dimensional space, and it generally deviates from this space during learning as shown in Fig. 6e.

As with the Inference model, a separate set of initial parameters 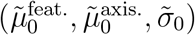 is used when the previous block was C2.

#### Structural Model

Our structural model parameterized control as a beta distribution over the space of low-dimensional task embeddings mapped onto the vertices of the underlying parallelogram (in this case a square), much like strong structural composition. Crucially, however, this model doesn’t enforce a strong prior over the vertices (prior is fit using free parameters, described below). In this case exploration takes place over the whole range of values between 0 and 1. To avoid slow learning dynamics around 0.5 which occur using the update rule in Eq. 64, we use an asymmetric learning rule. We also update each dimension of control with individual learning rates, *η*^feat.^ and *η*^resp.^

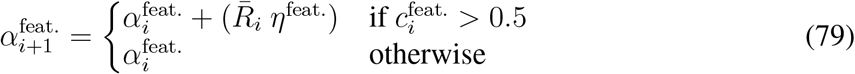

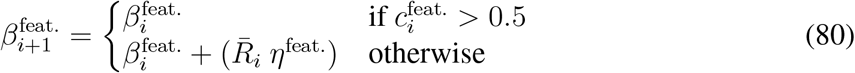

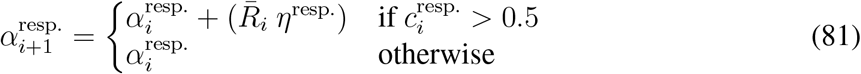

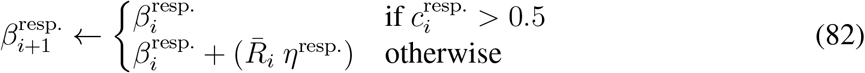

where 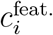 and 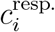 are the dimensions of control vector **c** that correspond to the feature and response axis dimensions respectively. Again we emphasize that the feature and response dimensions in the control subspace emerge naturally from pretraining on the compositional task set, and are not modeler defined. Here we simply label them with the corresponding subcomponent of the task set (feature or response axis).

After each trial, if the previous block was C2, both beta concentration parameters for the feature dimension are mixed toward a fixed regularization value *ρ*:

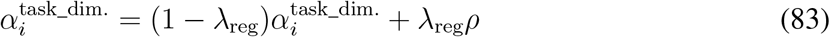

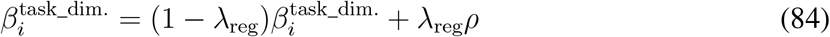

where task_dim. ∈ {feat., resp.}. We fix *ρ* = 0.1. At each trial we obtain 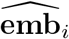 in the same manner as strong structural learning

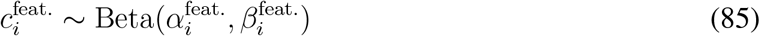

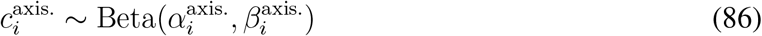

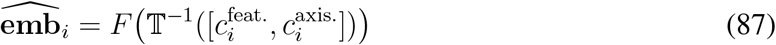

For prior parameters of each of the beta distributions, we fit mean 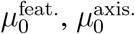 and concentration 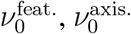 parameters and map these back to the appropriate *α* and *β* parameters.

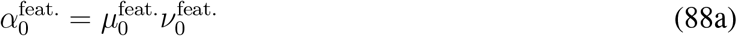

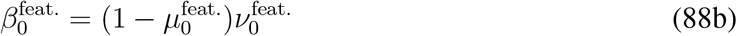

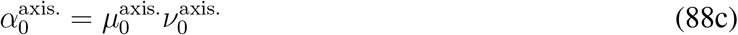

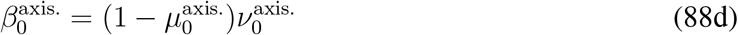

As with the previous models, when the previous block was C2 we use a separate set of prior parameters 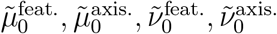.

#### Free Parameters

The free parameters for each model are given by

#### Parameter Fitting

The process of fitting parameters for each model proceeds in two phrases. First, we fit prior parameters using the distribution of animals’ initial actions in each block. Once these prior parameters are set, we then optimize the remain parameters to minimize mean squared error between animal and model task discovery trajectories.

Prior parameters were fit to minimize the negative log-likelihood of the animal’s first actions across blocks. Following the model descriptions above, in order to capture the possibility that animals use information in the sequence of blocks, for all models we fit two sets of prior parameters, one for C2 blocks (denoted *θ*_prior_) and one for S1/C1 blocks jointly (denoted 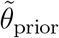). Let 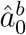 be the animals’ initial action in block *b* and let *B*_S1_, *B*_C1_, *B*_C2_ be the set of S1, C1, and C2 blocks respectively. The full prior loss is then

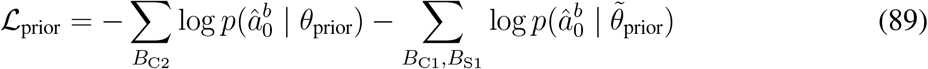

For the Modular and Structural models, the marginal is estimated by drawing *K* embedding samples from the prior distribution and averaging the resulting action probabilities:

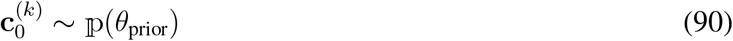

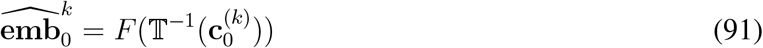

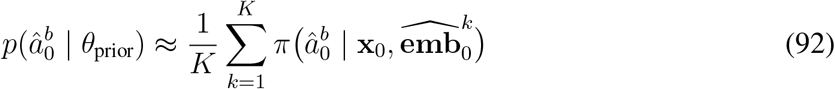

Where p represents the control distribution in each case i.e. a normal distribution for the Modular model and a Beta distribution for the structural model. For the Inference model, the marginal action probability at the first trial is computed exactly as a weighted sum over the three task embeddings, where the weights are given directly by the prior belief *b*_0_:

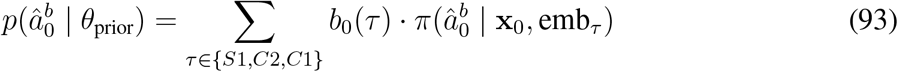

where *b*_0_(*τ*) = *w*_*τ*_ */ ∑*_*j*_ *w*_*j*_ are the normalized prior weights (or 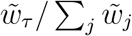 if the previous block was C2).

The second phase optimizes the remaining parameters by minimizing the mean squared error between the model and animal task discovery trajectories. The MSE is computed over smoothed reward trajectories using a moving average with window size 5:

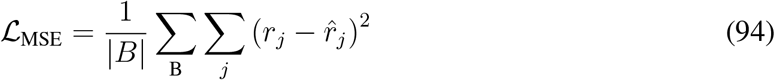

where *r*_*j*_ and 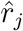 denotes the moving-average smoothed reward at trial *j* for the model and animal respectively. Both stages used CMA-ES (Covariance Matrix Adaptation Evolution Strategy [82]) as implemented in Optuna. Both stages were run for 250 iterations. Cross-validation used stratified 5-fold splitting (via scikit-learn’s `StratifiedKFold`), stratified by block rule to ensure balanced representation of S1, C2, and C1 blocks across folds. All evaluations and analyses in the main text are the result of running fitted parameters on blocks held out from the fitting procedure. During fitting we let *K* = 10.

### Decoding Analyses

The decoding analyses follow the methods of Tafazoli et al. [29]. In general, the goal is to track the re-engagement of different computational subcomponents by training decoders to map from RNN hidden state to some variable (e.g. stimulus color category) in late trials of one block type (e.g. C2) where performance is stable. Then we test the decoder on early trials of alternative blocks (e.g. C1) to determine whether representations are re-used, and, importantly, if there is a trend in cross-decoding performance as the models re-engage computations as the block progresses. To balance training distributions for decoder we separate trials by stimulus congruency, which describes whether stimulus shape and color features map to the same or different categories. There are four stimulus congruency classes in total:

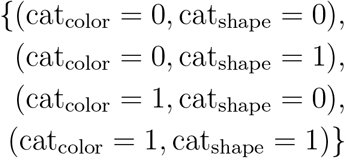

We trained logistic regression decoders for each within-trial time step on stimulus color category, stimulus shape category, and response axis direction. Again, following the approach in [29], for each decoding target we first pool eligible trials into training and testing distributions. We then train and test decoders on a small sample from these distributions, repeat this over many different decoder iterations, and average across decoders to obtain overall cross decoding performance. For each decoding target we trained 50 individual decoder for each of the 5 randomly initialized RNN models, resulting in 250 total decoders per trial time step.

#### Color Decoder

Color decoders were trained to decode the color category of the stimulus from RNN hidden states. Only the final correct 75 trials of the C2 block were eligible for inclusion in the training set. For each decoder, we drew four trials from each of the congruency classes. The test distribution was built by taking sliding windows (window size of 40 trials with a slide of 5 trials) from the first 110 trials of C1 blocks. For each window, 10 trials from each of the four congruency classes is selected and are used to evaluate cross decoding accuracy.

#### Shape Decoder

Shape decoders were trained to decode the shape category of the stimulus from RNN hidden states. Only the final 75 trials of the S1 block were eligible for inclusion in the training set. For each decoder, we drew four trials from each of the congruency classes. These training set were also balanced to include an even number of correct and incorrect trials. The distribution of test trials and the methods of sampling was the same as in Color Decoder.

#### Response Axis Decoder

Response axis decoders were trained to decode the direction of the model response from RNN hidden states. The final 75 trials of the S1 block were eligible for inclusion in the training set. In this case, for each decoder, we drew 16 trials which were balanced across trial correctness and direction (4 correct trials where model responded with direction 0, 4 incorrect trials where model responded with direction 0, 4 correct trials where model responded with direction 1, and 4 incorrect trials where model responded with direction 1). Like above 40 test trials are drawn from each window in the C1 blocks. In this case these trials are balanced in the same way in the training distribution.

For each decoder we subsampled 5% of RNN hidden neurons to train decoders. Decoders were implemented with scikit-learn’s `LogisticRegression` with the `liblinear` solver with L2 regularization (C=10.0). A null distribution was computed at each iteration by evaluating the trained decoder on permuted test labels. Decoding accuracy curves plotted in the main text are cross-decoding performance averaged over all decoders for each window of trials.

Cross decoding performance for neural data was taken directly from [29]. In that work, they analyzed trends in cross decoding across different sequences of blocks. We averaged across sequence types to obtain cross decoding on the whole dataset which we then compared to our models.

#### Decoder Trends Statistical Test

The cross decoding described above is primarily designed to examine whether or not there are trends in how models recruit compositional computations as a block progresses. To test the statistical significance of these trends, we again followed the approach in [29]. We used a trend test with cluster-mass correction applied to the sequence of decoding accuracies across trial windows and at each individual timestep within a trial. First, for a timestep to be considered for testing the difference between cross decoding performance at first and last trial window had to exceed an absolute value of 0.01. At each timestep *t* within a trial, we computed the Sen’s slope i.e. the median of all pairwise slopes of decoding performance of trial windows:

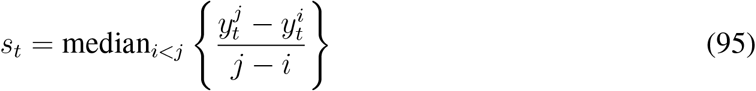

where *i* and *j* index the window of trials within a block and 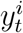 is the decoder test performances for trial window *i* at within trial timestep *t*. A null distribution of Sen slopes was generated by permuting the order of trial windows 250 times and recomputing the Sen’s slope at each timestep for each permutation. Significance was assessed using a two-tailed cluster-mass correction. Cluster masses were defined as the sum of the Sen’s slopes over contiguous time steps where computed slopes exceeded 95% of the null distribution. The p-values for each cluster was calculated as the proportion of permuted trial windows which resulted in cluster masses that were greater than those observed in the true cross decoding accuracies.

## Data and Code Availability

Code and pre-trained models are available at https://doi.org/10.5281/zenodo.21334463. All behavioral and neural data from the experiments of Tafazoli et al. can be found https://doi.org/10.6084/m9.figshare.30276238.v1.

## Acknowledgments

R.R. and P.L. were supported by the Gatsby Charitable Foundation (awards GAT3850 and GAT4058). A.P. was supported by grant 10.004.051 from the Swiss National Science Foundation. We’d also like to thank Charles Findling for helpful discussions, especially early on in the project, Jacob Bakermans for very constructive notes of early drafts of the manuscript and excellent discussions throughout, and Yedi Zhang for many helpful suggestions and references on the theoretical side of the project.

