## Supplementary material for "Structural Composition Enables Very Fast Learning": Mathematical Supplement

#### 1 Solution Space in Compositional Feedforward Networks

Suppose we have a teacher network that generates data according to

$$\mathbf{y}_\tau = \sum_{m=1}^{M} G^*_{\tau,m}\mathbf{W}^*_m\mathbf{x} \tag{S.1}$$

where $G^*_{\tau,m}$ is a scalar element of $\mathbf{G}^* \in \mathbb{R}^{\mathcal{T}\times M}$, $\mathbf{W}^*_m \in \mathbb{R}^{o\times d}$, $\mathbf{x} \in \mathbb{R}^d$, $\mathbf{y}_\tau \in \mathbb{R}^o$, $\mathcal{T}$ is the total number of tasks, and $M$ is the total number of linear modules in the teacher network. We'll assume that $\mathcal{T} \geq M$, each task uses at least one teacher module (no $\mathbf{0}$ rows in $\mathbf{G}^*$), and each module is used by at least one task (no $\mathbf{0}$ columns in $\mathbf{G}^*$). The teacher weights $\mathbf{W}^*_m$ are drawn from an independent normal distribution with mean 0 and variance $\sigma^2_m$, and the input is whitened,

$$(\mathbf{W}^*_m)_{kl} \sim \mathcal{N}(0,\sigma^2) \tag{S.2a}$$

$$\mathbf{x} \sim \mathcal{N}(\mathbf{0}_d, \mathbb{I}_d) \tag{S.2b}$$

where $d$ is large.

We train a student network on the dataset $\{\mathbf{x}^j, \mathbf{y}^j_\tau\}$ generated by the teacher. The student has the same structure as the teacher; the only difference is that it has $K$ modules instead of $M$,

$$\hat{\mathbf{y}}^j_\tau = \sum_{k=1}^{K} G_{\tau,k}\mathbf{W}_k\mathbf{x}^j\,. \tag{S.3}$$

We'll assume also that $\mathcal{T} \geq K \geq M$. The student updates $\mathbf{G}$ and $\mathbf{W}_k$ via gradient descent to minimize the mean squared error loss averaged over tasks,

$$L = \frac{1}{2\mathcal{T}}\sum_{\tau=1}^{\mathcal{T}} \left\langle ||\mathbf{y}_\tau - \hat{\mathbf{y}}_\tau||^2 \right\rangle \tag{S.4}$$

where the angle brackets indicate an average over the data, $\{\mathbf{x}^j, \mathbf{y}^j_\tau\}$.
In the limit of small learning rate we can write the update equations for $\mathbf{W}_k$ and $\mathbf{G}$ in terms of gradient flow,

$$\frac{d}{dt}\mathbf{W}_k = \sum_{\tau=1}^{\mathcal{T}} G_{\tau,k}\left\langle(\mathbf{y}_\tau - \hat{\mathbf{y}}_\tau)\mathbf{x}^\intercal\right\rangle \tag{S.5a}$$

$$\frac{d}{dt}G_{\tau,k} = \left\langle(\mathbf{y}_\tau - \hat{\mathbf{y}}_\tau)^\intercal(\mathbf{W}_k\mathbf{x})\right\rangle \tag{S.5b}$$

Explicitly performing the averages, and expressing $\mathbf{y}$ and $\hat{\mathbf{y}}$ in terms of $\mathbf{x}$ via Eqs. (S.1) and (S.3), respectively, we find, after a small amount of algebra, that the gradient flow equations are

$$\frac{dW_k}{dt} = \sum_{\tau=1}^{\mathcal{T}} G_{\tau,k}\left(\sum_{m=1}^{M} G^*_{\tau,m}\mathbf{W}^*_m - \sum_{l=1}^{K} G_{\tau,l}\mathbf{W}_l\right)\mathbf{\Sigma}_{xx} \tag{S.6a}$$

$$\frac{dG_{\tau,k}}{dt} = \text{tr}\left(\mathbf{W}_k^{\intercal}\left(\sum_{m=1}^{M} G^*_{\tau,m}\mathbf{W}^*_m - \sum_{l=1}^{K} G_{\tau,l}\mathbf{W}_l\right)\mathbf{\Sigma}_{xx}\right) \tag{S.6b}$$

where

$$\mathbf{\Sigma}_{xx} \equiv \langle \mathbf{x}^{\intercal}\mathbf{x} \rangle \,. \tag{S.7}$$

In our analysis we use a white covariance matrix, $\mathbf{\Sigma}_{xx} = \mathbb{I}$. We allow the covariance matrix to be arbitrary for generality.

To make contact with previous work on linear networks, we make the definitions

$$(\mathbf{g}_k)_\tau \equiv G_{\tau,k} \qquad (\mathbf{g}^*_k)_\tau \equiv G^*_{\tau,k} \tag{S.8a}$$
$$(\mathbf{w}_k)_{(ij)} \equiv (W_k)_{ij} \qquad (\mathbf{w}^*_k)_{(ij)} \equiv (W^*_k)_{ij} \tag{S.8b}$$
$$(\mathbf{\Sigma})_{ij,kl} \equiv \delta_{ik}(\mathbf{\Sigma}_{xx})_{jl} \,. \tag{S.8c}$$

Equation (S.8a) defines a vector with components corresponding to task, $\tau$; Eq. (S.8b) defines a vector with compound index $(ij)$; and Eq. (S.8c) extends the covariance matrix via a Kronecker product, but we used indices to avoid confusion. With these definitions, Eq. (S.6) becomes

$$\frac{d\mathbf{w}_k}{dt} = \mathbf{g}_k \cdot \left(\sum_{m=1}^{M} \mathbf{g}^*_m\mathbf{w}^*_m - \sum_{l=1}^{K} \mathbf{g}_l\mathbf{w}_l\right) \cdot \mathbf{\Sigma} \tag{S.9a}$$
$$\frac{d\mathbf{g}_k}{dt} = \left(\sum_{m=1}^{M} \mathbf{g}^*_m\mathbf{w}^*_m - \sum_{l=1}^{K} \mathbf{g}_l\mathbf{w}_l\right) \cdot \mathbf{\Sigma} \cdot \mathbf{w}_k \,. \tag{S.9b}$$

These are exactly the equations for a two layer linear network. When starting from small weights, then at the end of learning [1],

$$\mathbf{g}_k = \sum_{\mu}^{r} a_{k\mu}\mathbf{u}^*_\mu \tag{S.10}$$

where the $\mathbf{u}^*_\mu$ are the left singular values associated with the teacher $\mathbf{G}$,

$$\sum_{m=1}^{M} \mathbf{g}^*_m\mathbf{g}^*_m \cdot \mathbf{u}^*_\mu = s^2_\mu\mathbf{u}^*_\mu \,, \tag{S.11}$$

and the sum in (S.10) includes only values of $\mu$ for which $s_\mu \neq 0$; i.e., there are $r$ terms where $r$ is the rank of $\mathbf{G}^*$.

It is convenient to return to our original notation (we introduced new notation solely to make contact with linear networks). First, Eq. (S.11) implies that the singular value decomposition of $\mathbf{G}^*$ is

$$\mathbf{G}^* = \mathbf{U}^*\mathbf{S}^*\mathbf{V}^{*\intercal} \tag{S.12}$$

where the rows of $\mathbf{U}$ correspond to the vectors in $\mathbf{u}^*_\mu$ that appear in Eq. (S.10) and (S.11), $\mathbf{S}$ is a diagonal matrix with the singular values, $s_\mu$ along the diagonal, and $\mathbf{V}$ is a matrix containing the right singular vectors. Second, Eq. (S.10) can be written

$$\mathbf{G} = \mathbf{U}^*\mathbf{A} \tag{S.13}$$

where $\mathbf{A}$ is the transpose of the matrix $a_{k\mu}$ that appeared in Eq. (S.10).
To determine how the left singular vectors of $\mathbf{G}$ are related to those of $\mathbf{G}^*$, first write $\mathbf{G}$ as

$$\mathbf{G} = \mathbf{U}\mathbf{S}\mathbf{V}^{\mathsf{T}}\,. \tag{S.14}$$

Combining this with Eq. (S.13) then yields

$$\mathbf{U} = \mathbf{U}^*\mathbf{Q} \tag{S.15}$$

where

$$\mathbf{Q} \equiv \mathbf{A}\mathbf{V}\mathbf{S}^{-1}\,. \tag{S.16}$$

Using the orthogonality of the singular vectors, we have

$$\mathbf{U}^{\mathsf{T}}\mathbf{U} = \mathbf{Q}^{\mathsf{T}}\mathbf{U}^{*\mathsf{T}}\mathbf{U}\mathbf{Q} = \mathbf{Q}^{\mathsf{T}}\mathbf{Q} = \mathbb{I}\,. \tag{S.17}$$

Consequently, $\mathbf{Q}$ is a rotation matrix (with a possible reflection). This immediately implies that

$$\mathrm{rank}(\mathbf{G}) = \mathrm{rank}(\mathbf{G}^*)\,. \tag{S.18}$$

Now consider the centered matrices $\tilde{G}$, $\tilde{G}^*$, where each column has 0 mean. We can express these matrices as a product of the center mapping $\mathbf{C} = (I - \frac{1}{\mathcal{T}}\mathbf{1}_{\mathcal{T}}\mathbf{1}_{\mathcal{T}}^{\mathsf{T}})$, where $\tilde{\mathbf{G}} = \mathbf{C}\mathbf{G}$ and $\tilde{\mathbf{G}}^* = \mathbf{C}\mathbf{G}^*$. This centering reduces the rank of the resulting matrix if $\mathbf{1}$ is in the column space. From (S.22) we know that $\mathbf{G}$ and $\mathbf{G}^*$ have the same column space, so centering will have the same effect on the rank in both cases and

$$\mathrm{rank}(\tilde{\mathbf{G}}^*) = \mathrm{rank}(\tilde{\mathbf{G}}) \equiv \tilde{r} \tag{S.19}$$

Further, from (S.13) we have that

$$\tilde{\mathbf{G}} = \mathbf{C}\mathbf{G} = \mathbf{C}\mathbf{U}^*\mathbf{A} \tag{S.20}$$

$$\tilde{\mathbf{U}}^* = \mathbf{C}\mathbf{U} \tag{S.21}$$

where $\tilde{\mathbf{U}}^*$ is a $\mathcal{T} \times \tilde{r}$ matrix collecting the remaining independent orthonormal columns of $\mathbf{U}^*$ after centering. In this case, the apply the same argument as (S.13)-(S.22) to obtain

$$\tilde{\mathbf{U}} = \tilde{\mathbf{U}}^*\tilde{\mathbf{Q}} \tag{S.22}$$

where $\tilde{\mathbf{Q}}$ is an $\tilde{r} \times \tilde{r}$ rotation matrix.

### 2 Geometry of Learned Representations for Binary $\mathbf{G}^*$

In our formulation its useful to think of the first $r$ components of the rows of the left singular vectors, denoted $\mathbf{u}^*[\tau, :r] \equiv \mathbf{u}^*_\tau$, as the coordinates for task $\tau$ in $r$-dimensional space. If $\mathbf{G}^*$ is made up of any real values, the actual realization of the geometry formed by these coordinates can be very general. However, in the experimental literature and in many real-world settings, the primary concern is often whether or not a module should be included in a task, rather than its precise weighting. In these cases the gating matrix $G^*$ is binary and the set of $\mathbf{u}^*_\tau$ form a subset of the vertices of a parallelepiped with at most $2^r$ vertices.

To see this consider the truncated SVD of binary matrix $\mathbf{B}$ or rank $r$.

$$\mathbf{B} = \mathbf{USV}^\intercal \tag{S.23}$$

where $\mathbf{B} \in \{0,1\}^{n \times m}$. By definition, the rows of $\mathbf{B}$ lie on the vertices of a hypercube in $m$-dimensional space. We can then invert the SVD mapping and write

$$\mathbf{BVS^{-1}_{m,r}} = \mathbf{U} \tag{S.24}$$

where $\mathbf{S^{-1}}$ is an $m \times r$ matrix with the scaler inverse of the singular values along the diagonal. The map on the left hand side of S.15 itself is enough to develop a strong intuition about the geometries that should emerge. We begin with a subset of the vertices of a $m$-dimensional hypercube in the rows of $\mathbf{B}$. This is then rotated through $\mathbf{V}$, an operation that preserves all the geometric properties of the original hypercube. $\mathbf{S^{-1}}$ then annihilates $m - r$ dimensions and scales the remaining dimensions. This results is a subset of the vertices of an $r$-dimensional hypercube with scaled directions i.e. a $r$-dimensional parallelepiped which $2^r$ vertices.

Generically this is true for any linear map $A : \mathbb{R}^m \to \mathbb{R}^r$ acting on $\mathbf{B}$ given that the columns of $\mathbf{A}$ are linearly independent and $r < m$. The set of rows of $\mathcal{Z} = \mathbf{BA}$ are given by some linear combination of the rows of $\mathbf{A}$.

$$\mathcal{Z} = \left\{ \sum_{i=1}^{m} \beta_i A_i^\intercal \;\mid\; \beta \in \{0,1\}^m \right\}$$

Linear independence in the columns of $\mathbf{A}$ means combinations of exactly the first $r$ rows are sufficient to capture this set.

$$\mathcal{Z} = \left\{ \sum_{i=1}^{r} \beta_i A_i^\intercal \;\mid\; \beta \in \{0,1\}^r \right\}$$

Then we have a full rank transformation for of a subset of $r$-dimensional hypercube vertices which results in a subset of parallelepiped vertices.

### Supplemental References

[1] Arthur Jacot, François Ged, Berfin Şimşek, Clément Hongler, and Franck Gabriel. Saddle-to-saddle dynamics in deep linear networks: Small initialization training, symmetry, and sparsity. *arXiv preprint arXiv:2106.15933*, 2021.
