## Supplementary Figures for "Structural Composition Enables Very Fast Learning"

### Supplementary Information

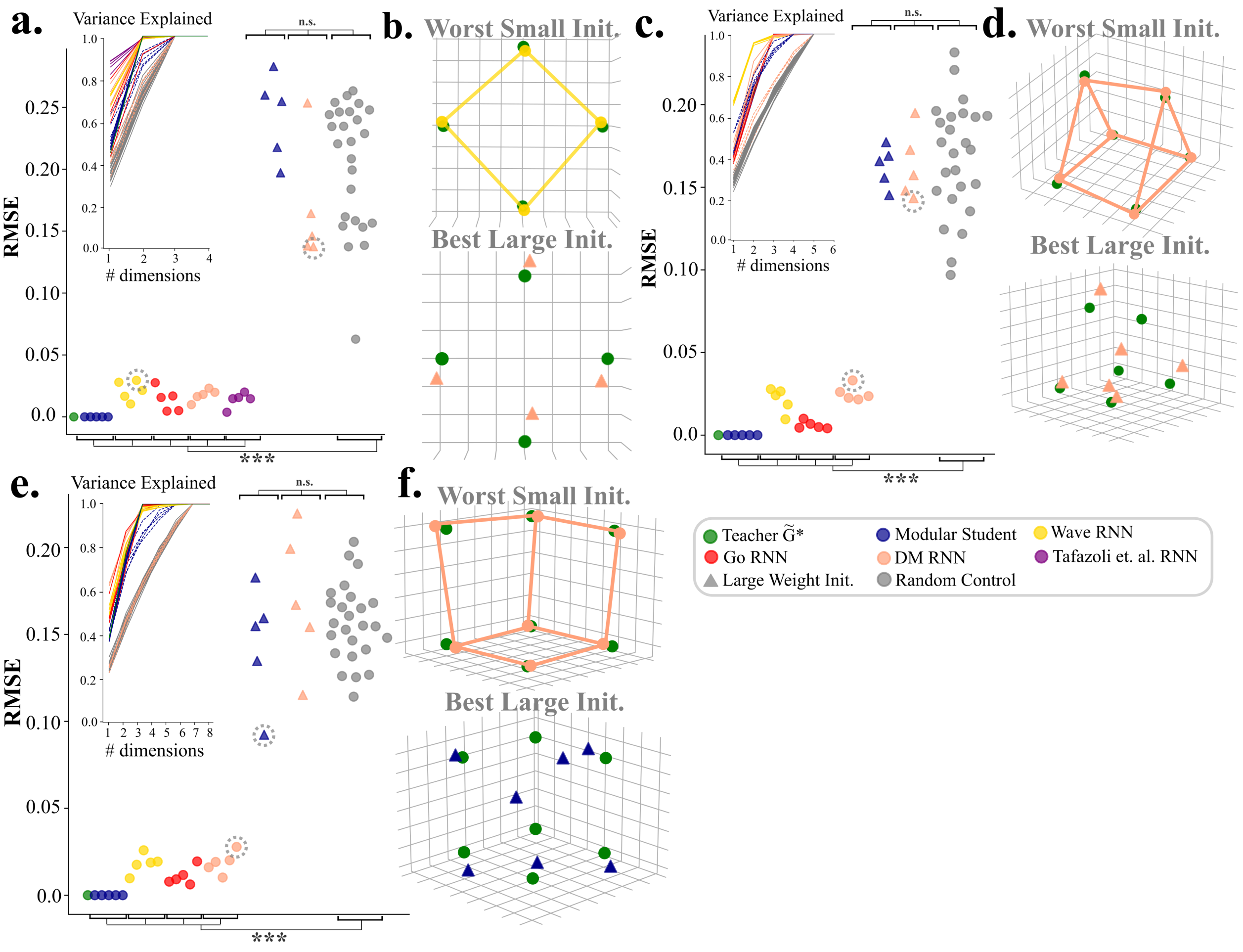


**Supplementary Figure 1: Task set structure is learned in a variety of compositional structures. a.-b.** Same as Fig. 3d-e. main text for 2-D task set configuration (see Methods, Task Classes). **c.-d.** Same as Fig. 3d-e. main text for 3-D (6) task set configuration (see Methods, Task Classes). **e.-f.** Same as Fig. 3d-e. main text for 3-D task set configuration with one task held out of training.

Here we test three additional kinds of compositional structure to provide additional evidence that structure in trained embedding weights $\widetilde{\mathbf{W}}_{\text{emb}}$ are well approximated by Eqs. **9** in the main text. The measures that are plotted here follow Fig. 3 in the main text.

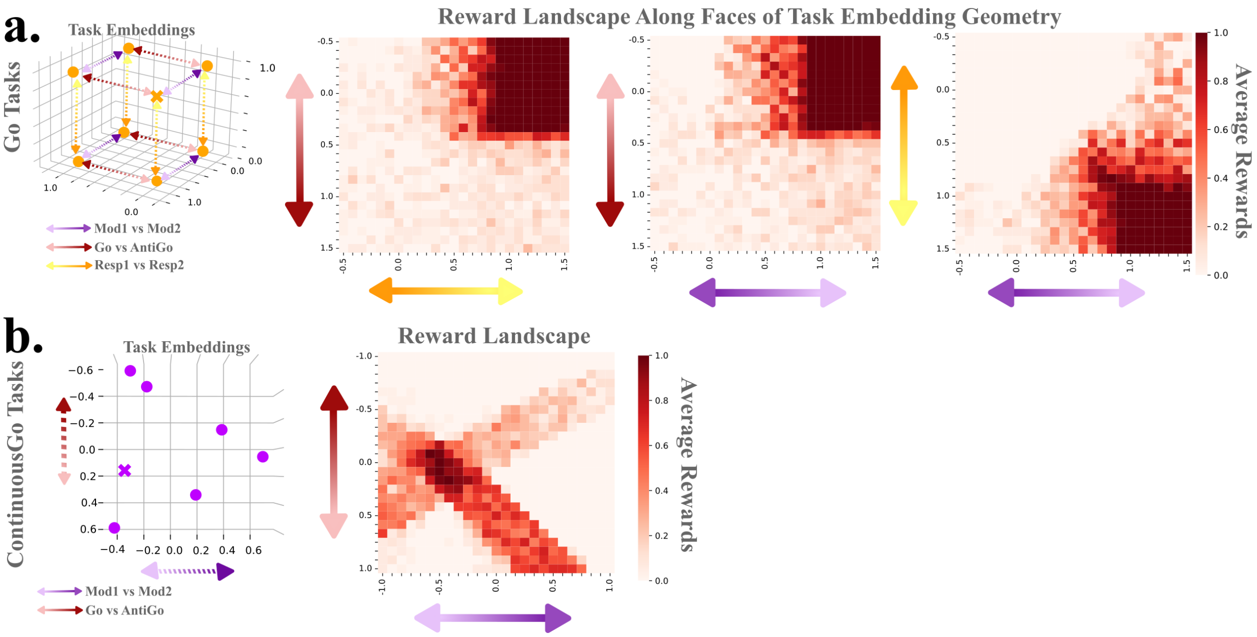


**Supplementary Figure 2: Reward Landscape for Binary vs. Continuous Task Compositions. a.** Left most inset shows the task embedding cube along with the task missing vertex that leads to high reward for the heldout Mod1-Go-Resp1 task (marked with the x). Right three subplots show reward landscapes for the heldout Mod1-Go-Resp1 task evaluated on the three faces of the underlying task embedding cube that intersect the correct embedding. Arrows labeling the axes show which task set components lie along each of the axes. **b.** Same as **a.** for the ContinuousGo task set. The reward landscape is plotted on the plane that defines the task embedding space.

Here we show reward landscapes for the Mod1-Go-Resp1 when held out of training (Supplementary Fig. 2a.) and for the unseen task in the ContinuousGo (Supplementary Fig. 2b.) task evaluated in the Fig. 4g. For the Mod1-Go-Resp1 task, reward landscapes are calculated on the faces of the unit cube that structures the transformed control space (see Methods, Compositional Learning from Reinforcement). Specifically, we evaluate a 25 by 25 grid of points $(\epsilon_1, \epsilon_2)$ ranging between (-0.5, 1.5) on each dimension. We set the $\bar{\mathbf{c}} = (c, \epsilon_1, \epsilon_2)$ where $c$ is set to 0 or 1 to match the plane where the correct Mod1-Go-Resp1 resides. $\bar{\mathbf{c}}$ is mapped back into the full embedding space (again, see Methods) and the RNN is evaluated on 100 trials of Mod1-Go-Resp1 using that embedding. Heatmap shows average reward obtained over those 100 trials. This process is repeated for $c$ in the 2nd and 3rd dimensions of $\bar{\mathbf{c}}$ to obtain all three performance heatmaps. The analysis is the same in Supplementary Fig. 3b. for the ContinuousGo task, only in this case $\widetilde{\mathbf{W}}_{\text{emb}}$ is rank 2 so we evaluate $\bar{\mathbf{c}} = (\epsilon_1, \epsilon_2)$ over the same grid resolution.

We plot these heatmaps to show that the compositional structure of the task set can affect the underlying reward landscape in significant ways. In binary tasks (Go Task set) there is a large region of space that lead to high reward. This region appears to fully occupy the corner of the underlying cube where the correct Mod1-Go-Resp1 embedding resides. By contrast in the novel ContinuousGo task, highly rewarding regions tend to be smaller and are much more tightly centered around the correct embedding itself. Speculatively, this may be due to effects of the non-linearity when the network must implement either one set of dynamics or the other (as in the Go task set), as opposed to a smooth interpolation between mappings (as in the ContinuousGo task set).

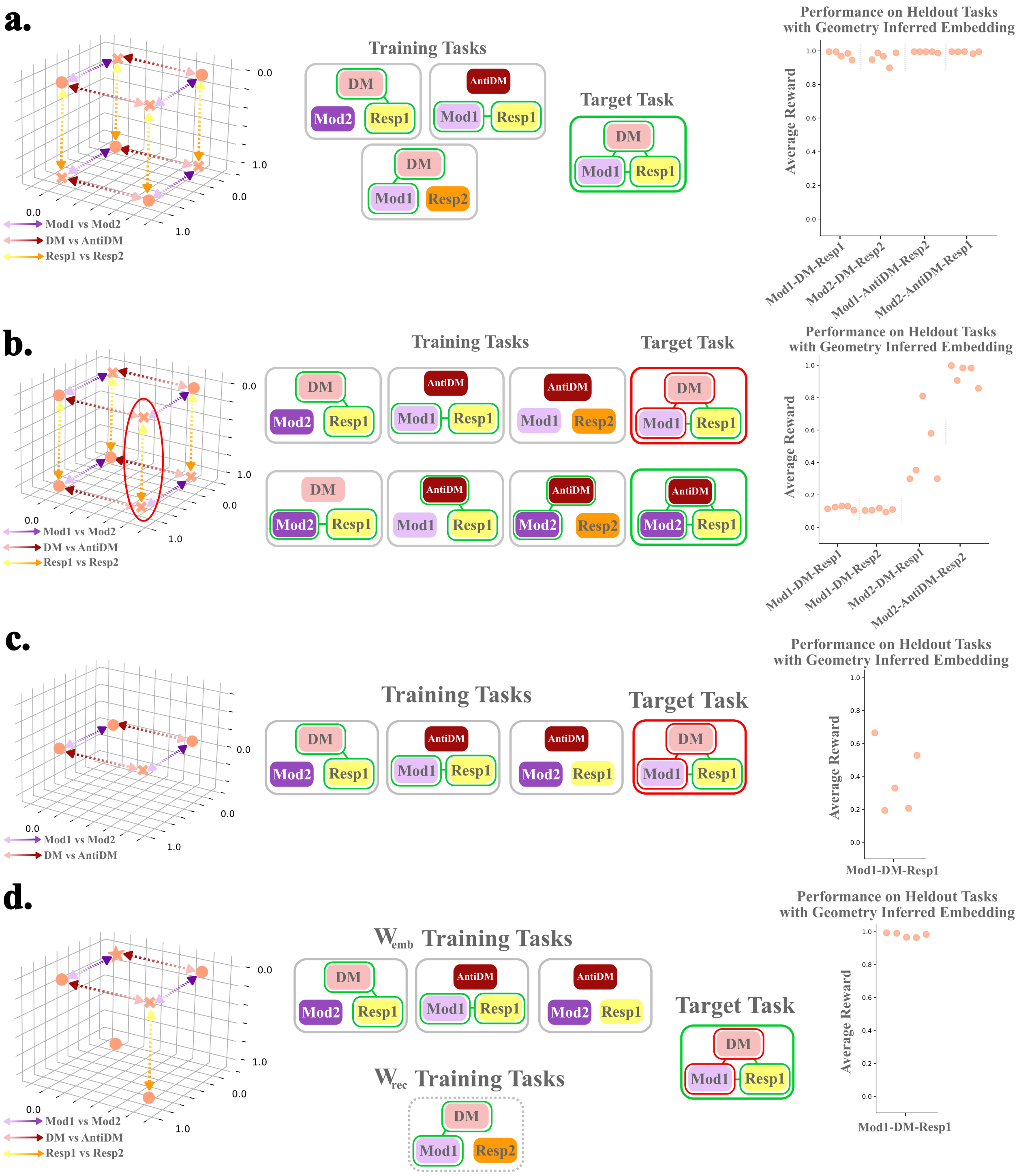


**Supplementary Figure 3: Generalization with multiple tasks held out. a.** Task embeddings and performance for 3-dim DM task set with 4 tasks held out of training. Note that all held out tasks (marked by x's) are anti-podal i.e. each is adjacent to three tasks in the training set (dots). This means that the RNN has all the connected components need to generalize to all held out tasks. An example is shown in the middle inset where Mod1-DM-Resp2; Mod1-AntiDM-Resp1; Mod2-DM-Resp1 in the training set allow generalization to the held out Mod1-DM-Resp1 task. Each dots in averaged reward is performance for a model with a different random initialization evaluated with the proper vertex of the embedding cube across many trials of the heldout task. The result is high performance across all held out tasks. **b.** Same as **a.** but for a configuration of held out tasks that is missing a connected component. Examples of a sufficient and insufficient set of training tasks are shown in the middle insets. Only the task with all required connected components (Mod2-AntiDM-Resp2, with 3 adjacent training tasks) shows high performance. **c.** Same **a.** for the 2-dim. DM task set with one task held out of training. **d.** Same **c.** but recurrent weight $\mathbf{W}_{rec}$ are trained on the same set of tasks as in **a.** and embedding weights $\mathbf{W}_{emb}$ trained from scratch only on tasks from **c.**. In this case having all recurrent components results in high performance on the held out task.

In the main text for clarity and simplicity we only considered task sets with a single task held out for testing. A natural follow up question is: what is the minimal number of training tasks required to support performance in the task set overall? For structural composition, training tasks need to support the underlying geometry of the task set. To construct an $r$-dimensional subspace containing specific coordinates, $r + 1$ vectors are needed: one to fix a point in space and $r$ linearly independent vectors to define the directions. So for our 3-binary axis DM task set, we require four tasks, and not all of them can have the same component along one of the axes (e.g. if all four tasks involved Mod1, the task embeddings would simply form a plane capturing one of the faces of the underlying 3-d parallelogram). Given that this requirement is satisfied, the task embedding space in principle has all of the geometric properties needed to constrain the task set for weak structural composition. For strong structural composition, there is an additional geometric consideration. If the system is relying strictly on the geometry of embeddings, then the system has to deal with the ambiguity of how to complete parallelograms with missing vertices. For example, when a task is held out of the 2-dim structure shown in Supplementary Fig. 1a. the embeddings form an equilateral triangle, and the underlying parallelogram can be completed using three different points. It is simple to keep track of the adjacency to tasks by simply keep track of which task use which modules. With this knowledge the requirement of strong and weak structural composition are the same.

Empirically, we also find that for a nonlinear recurrent network to perform held out tasks, its components also have to "connect" during training. Roughly, this means that for the network to generalize to a held out task (e.g. Mod1-DM-Resp1) without any further tuning of recurrent weights, all pairwise components of the held out task must be present in the training set (e.g. Mod1-DM-Resp2; Mod1-AntiDM-Resp1; Mod2-DM-Resp1). This property was also shown in simple compositional tasks with nonlinear feedforward networks [1]. For our tasks, it is still possible to train on only 4 of the 8 tasks and have the model perform well on the entire task set, however these tasks have to chosen correctly. An example of such a successful configuration is shown in Supplementary Fig. 3a. We also show an example with 4 tasks improperly chosen (Supplementary Fig. 3b.), and demonstrate that those tasks in the task set which are "connected" perform well while those missing a connected component exhibit low performance. This requirement also explains why tasks with 2 binary axes (e.g. DM vs. AntiDM, Mod1 vs. Mod2; four total tasks whose embeddings form a square, see Supplementary Fig. 1a.) struggle to support generalize to unseen tasks even though embedding weights exhibit our desired geometric properties. In this case, training on three of the four tasks will always leave a disconnected component in the held out task, resulting in poor generalization (Supplementary Fig. 3c.). We emphasize that this is primarily a limitation in the recurrent dynamics of our model RNNs that results from training on an insufficiently broad set of tasks. Suppose alternatively we start with an RNN whose recurrent weights have been trained on the set of 4 tasks from 3-dim task that supported generalization in Supplementary Fig. 3a. If we now freeze recurrent weights and simply retrain a new $\mathbf{W}_{\text{emb}}$ on 3 out of the 4 tasks then the network will generalize successfully. This broader range of pre-existing skill may better capture the state of knowledge of animals in experiments. Through a combination of evolution and experience animals likely come into any situation with a very vast repertoire of abilities already available (broad RNN pretraining). More experience intensive shaping may indicate which abilities are relevant to the current context (task set specific $\mathbf{W}_{\text{emb}}$ training). Once these are known then animals can quickly shift between them to adapt to moment by moment changes in the demands of the environment (compositional learning).

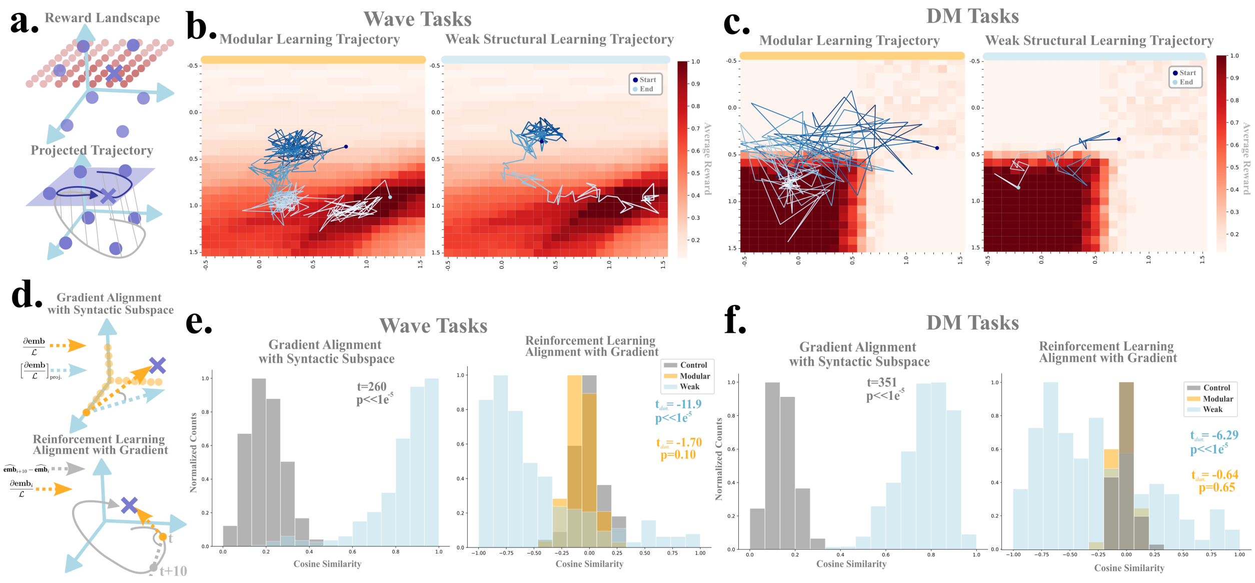


**Supplementary Figure 4: Weak structural learning approximates gradient descent using reinforcement. a.** Illustration of projecting weak structural learning on to reward landscape. Top shows the evaluation of the reward landscape across one face of the cube like in Supplementary Fig. 3a. Bottom shows projection of learning trajectory on the plane where reward landscape was evaluated. **b.** Example learning trajectories for a held out task in 3-dim wave task set. Left shows modular learning and right shows weak structural learning. **c.** Same as **b.** but for 3-dim DM task set **d.** Illustration of gradient alignment analyses. Top shows how we measure the gradient alignment with the structural embedding subspace encoded in $\widetilde{\mathbf{W}}_{\text{emb}}$. We take the gradient of the full embedding with respect to the true loss on the held out task. We then find the best projection of this gradient onto the relevant subspace and measure the cosine similarity between the projected and true gradient. We perform this analysis across a grid of points to measure how well the gradient across the full space aligns with the subspace. Bottom shows how we measure alignment between gradients updates from reinforcement. We take the gradient with respect to the full embedding at some trial $i$ in learning. Then we take the different between embedding at trial $i + 10$ and $i$ and measure the cosine similarity between this difference vector and the true gradient. Negative values on this measure indicate that learning is descending the gradient. **e.** Gradient alignment analysis in the wave task set. Left shows the distribution of the gradient alignment with the structural subspace. Right shows the distribution of alignment between gradients and updates from reinforcement for weak structural learning and modular learning. **f.** Same as **f.** but for the DM task set.

In the main text we showed that constraining learning to proceed within the relevant subspace of task embedding activity vastly speeds learning. This is in part because the dimensionality of the learning problem is reduced from the full embedding dimension (an arbitrary architectural choice) to the number of dimensions required to encode the patterns of module reuse across the task set (i.e. the rank of the $\mathbf{G}^*$). Here we show that full gradients of task embedding activity with respect to the true loss on the held out tasks largely reside within the $r$-dimensional subspace defined by $\widetilde{\mathbf{W}}_{\text{emb}}$. Hence, for weak structural learning updates from reinforcement constrained to this subspace have a relatively high probability of descending the gradient.

To start, we visualize the time course of learning for both modular and weak structural composition by plotting learning trajectories on a heatmap of reward. To do this we take a region of the task embedding space centered on one of the faces on the underlying cube of embeddings which includes the target task. We then take a grid of points in this plane and evaluate performance on the target task for each task embedding values in the grid just like in Supplementary Fig. 2a. This evaluation is illustrated in Supplementary Fig. 4a. top, where each red circle is an evaluation point and the intensity corresponds to the amount of reward received at that point. To show how learning explores this space, we take the full trajectory of embeddings vectors traversed during learning and project those vectors onto the plane (illustrated in Supplementary Fig. 4a. bottom). Representative runs of learning for the held out task in the 3-dim wave task set (Fig. 4d., main text) are plotted in

Supplementary Fig. 4b. Darker colors indicate early phases of learning while light colors indicate the end of learning. Here modular learning spends many trials exploring a suboptimal part of space before moving to find the highly rewarding region. By contrast, weak syntactic learning moves relatively smoothly after its initial phase of exploration to the rewarding region. Supplementary Fig. 4c. plots learning trajectories for the held out task in the DM task set (Fig. 4d., main texxt). For modular learning even though trajectories projected onto the place reside in the rewarding region, the model fails to find reward because of the large number of alternative dimensions it must explore. The model spends the majority of learning traversing these other dimensions before it finds a rewarding region close to the plane. Because weak structural composition is constrained, it need only explore 2 alternative dimensions once its enters the rewarding regions of the plane. This results in the relatively fast acquisition of the task after the initial phase of exploration.

The reduced dimensionality of weak syntactic composition improves the speed of learning. Importantly, however, these dimensions also define the *correct* subspace for learning, in the sense that the gradient of the full embedding with respect to the true loss largely resides in this subspace. To show this we again take a region of embedding space centered around trained task embeddings and now a 3-dimensional grid of points along the dimensions which define the structural subspace. In particular, we take the grid of points $\bar{\mathbf{c}} = (\epsilon_0, \epsilon_1, \epsilon_2)$ where $\epsilon_0, \epsilon_1, \epsilon_2$ each take on 25 evenly space values on the interval $(-0.5, 1.5)$. We then transform these grid points into the full embedding space (see Methods), and evaluate the loss on 100 trials of the held out task using the resulting embeddings. We can then calculate the gradient of the full embedding with respect to the loss at each point on the grid. For each of these gradients we then find the best projection onto the space defined by $\mathbf{W}_{\text{emb}}$. Finally, we compute the cosine similarity of the projected gradient with respect to the true gradient at each point on the grid. This procedure is illustrated in Supplementary Fig. 4d. top. The distribution of similarities for the wave task set is shown in Supplementary Fig. 4e. left. We find that in the region of space around trained task embeddings, the true gradient aligns strikingly well with the relevant subspace. Though reward functions across task classes differ in their details, in general an embedding that minimizes loss will attain high reward. Hence, changing the embedding in the direction of the negative gradient is the optimal update on any learning step. This result shows that the constraint of weak structural learning not only reduces the dimensionality of the learning problem, but confines updates to the same subspace where the majority of gradient information resides. This makes updates according to the approximate gradient much more likely even when learning only from trial-by-trial reinforcement. To see this, at each point in a learning trajectory $\widehat{\mathbf{emb}}_i$ we evaluate the full gradient of the embedding with respect to the true loss. We then take the point in the trajectory 10 trials afterwards, compute the vector $\widehat{\mathbf{emb}}_{i+10} - \widehat{\mathbf{emb}}_i$, and evaluate the cosine similarity between the result and the true gradient (Supplementary Fig. 4d. bottom). We do this for every learning episode. The distribution of cosine similarities between true gradients and reinforcement updates after ten trials for the wave tasks is shown is shown in Fig. 5e. right. First note that in modular learning, which freely explores the entire embedding space, the distribution of gradient alignments shows a very weak effect towards the negative gradient. By contrast, the gradient alignments for weak structural learning is very significantly shifted to negative values, meaning that a large proportion of updates approximately descend the gradient of the true loss. Again, this is true even though updates are made wholly from reinforcement on a trial-by-trial basis. The same results hold for an analysis of gradient alignments for the DM task class shown in Supplementary Fig. 4f. In all, these results demonstrate that by constraining learning to the relevant subspace, the dimensionality of the learning problem is reduced, but importantly it is reduced to a space which largely aligns with gradient updates of task embeddings. Hence, updates based on rewards have a relatively high likelihood of descending the gradient, resulting in much more efficient learning than the unconstrained, modular setting.

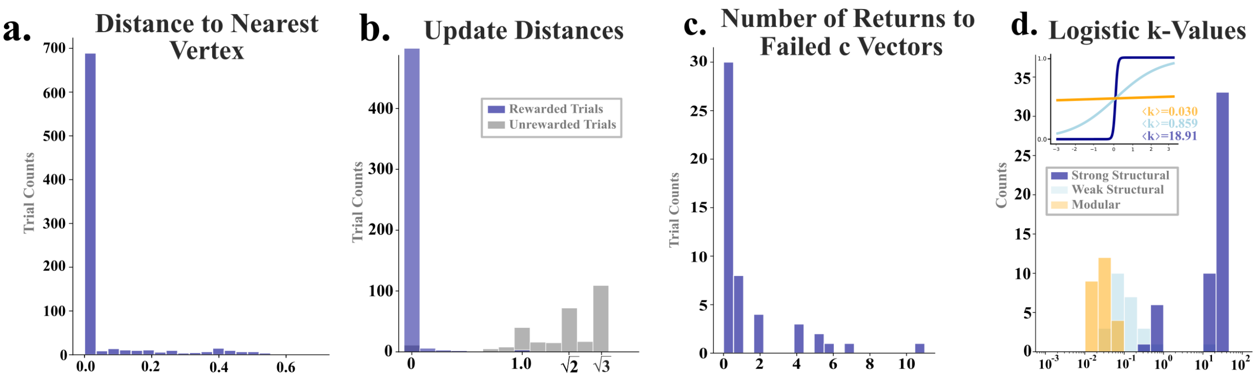


**Supplementary Figure 5: Strong structural learning implements hypothesis testing style learning. a.** Distribution of the distances of each vector **c** to the nearest vertex of the underlying cube during strong structural learning for the Mod1-Strong-Resp1 task. Includes every trial of 10 independent runs for each of the 5 different randomly initialized pretrained RNNs. **b.** Distribution of the updates distances after rewarded and unrewarded trials. **c.** Distribution of number of times the control vector **c** returns to a region that previously failed to collect reward. Here 30 of the 50 total learning runs contain no returns to failed regions. **d.** Distribution of k parameters for fitted logistic learning curves for each kind of learning. Inset plots a zero aligned curve with average k value for each kind of learning.

In Fig. 5. in the main text we outlined how strong structural learning implements a hypothesis testing style of learning. Here we extend that analysis by quantifying other relevant characteristics of learning. All analyses here are done on learning for the Mod1-Strong-Resp1 task. Intuitively, the strong prior on exploration in strong structural learning samples from points on the vertices of the cube. Supplementary Fig. 5a. quantifies this more precisely. It shows the distribution of distances between each vector **c** proposed during learning and the closest vertex. This distribution includes every trial over 50 total learning runs (10 runs among 5 pretrained RNNs initialized with different random seeds). Supplementary Fig. 5b. shows the distribution of update distances after rewarded and unrewarded trials across all learning. We find that reward confirms a hypothesis and results in **c** remaining at that embedding. By contrast, after an unrewarded trial the control vector moves to an entirely new vertex of the underlying cube (represented by distances peaked around 1.0, movement to an adjacent vertex, $\sqrt{2}$, a vertex diagonal on the face, and $\sqrt{3}$ an antipodal vertex). Supplementary Fig. 5c. show how often the control vector **c** returns to a region of space (a sphere radius 0.1) after failing to receive reward from that region on a previous trial. This measure how consistently strong structural learning eliminates a hypothesis after failing to receive reward. For 30 out of a total of 50 learning runs strong structural learning never returns to an unrewarded region. We showed in the main text that individual learning curves have sharp and sudden increases in performance. To quantify this across all learning runs and different learning types we fit logistic curves across each learning run individually. In fitting these curves the parameter $k$ controls the slope of learning. Very high values of $k$ are indicative of moments of insight. Distributions of $k$ values fit across all runs for each learning method are shown in Supplementary Fig. 5d. For strong structural composition, $k$ values tend to be an order of magnitude greater than those of weak structural composition. The inset shows a zero-centered curve plotted using the average $k$ value for each learning type. Strong structural learning exhibits a near step-wise increase in performance.

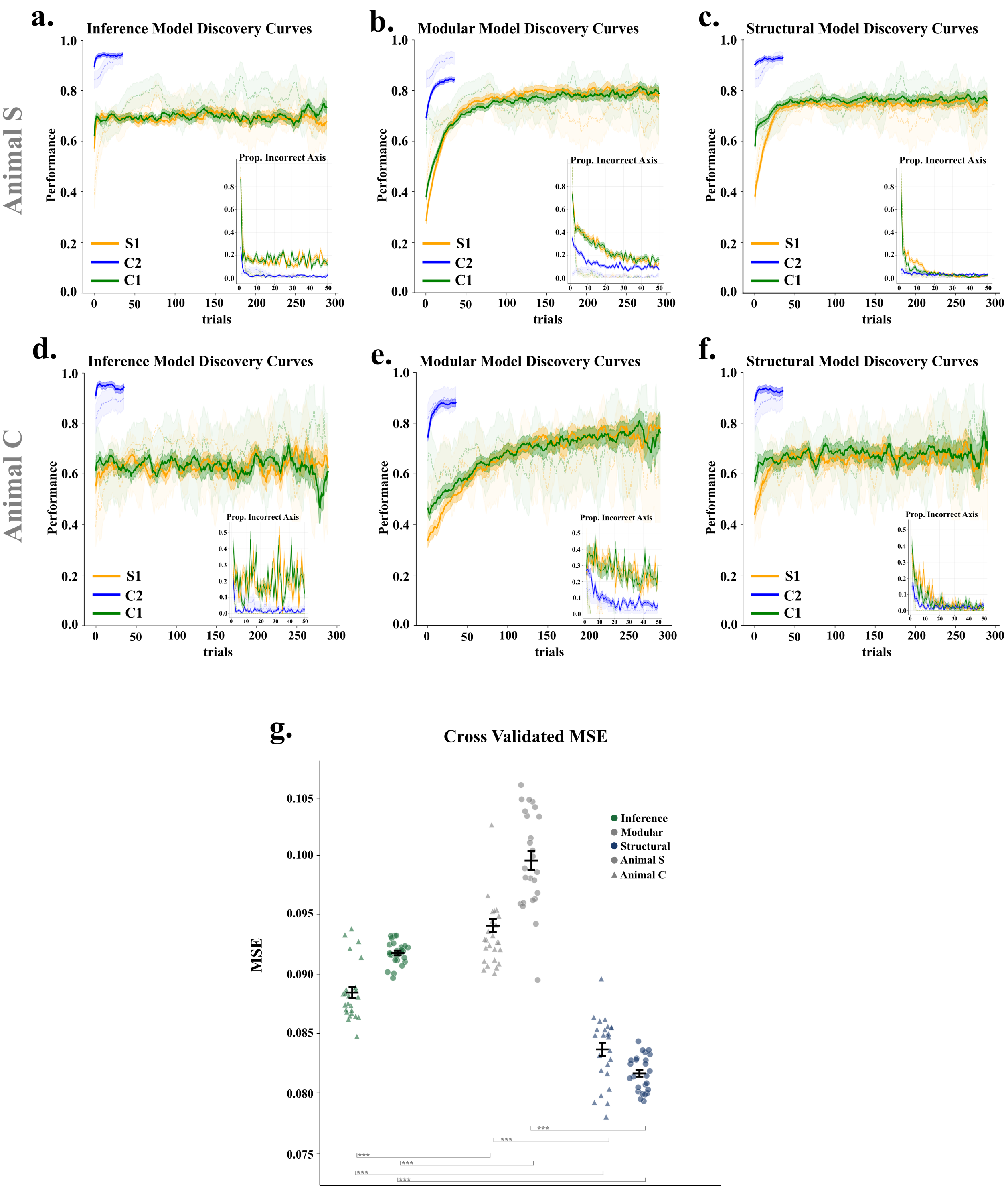


**Supplementary Figure 6: Model fits by individual animals. a.-c.** Same as the main text Fig. 6g.-i. except for data from monkey S only. **d.-f.** Same as the main text Fig. 6g.-i. except for data from monkey C only. **g.** Same as main text Fig. 6j. but for each animal individually.

### Supplementary References

[1] Simon Schug, Seijin Kobayashi, Yassir Akram, Maciej Wołczyk, Alexandra Proca, Johannes von Oswald, Razvan Pascanu, João Sacramento, and Angelika Steger. Discovering modular solutions that generalize compositionally, 2024.
